# Transcriptional regulation of neonatal neural stem cells is a determinant of social behavior

**DOI:** 10.1101/2021.11.12.468452

**Authors:** Takeshi Hiramoto, Shuken Boku, Gina Kang, Seiji Abe, Mariel Barbachan e Silva, Kenji Tanigaki, Masako Nagashima, Kenny Ye, Takahira Yamauchi, Tatyana V. Michurina, Pilib Ó Broin, Grigori Enikolopov, Noboru Hiroi

**Author notes:** SB, Department of Neuropsychiatry, Faculty of Life Sciences, Kumamoto University, Kumamoto, Japan SA Department of Hospital Pharmaceutics, School of Pharmacy, Showa University, Tokyo, Japan, TY Department of Psychiatry, Nara Medical University, Nara, Japan, MN Department of Palliative Medicine, Teikyo University School of Medicine, Tokyo, Japan. These two authors contributed equally to this work (Takeshi Hiramoto, Shuken Boku).

## Abstract

Rare gene variants confer a high level of penetrance to neurodevelopmental disorders, but their developmental origin and cellular substrates remain poorly understood. To address this limitation, we explored the role of *TBX1*, a gene encoded in a rare copy number variant, in cell and mouse models. Here, we report that neonatal *Tbx1* deficiency contributes to defective peripubertal social behavior and impairs the proliferation of neonatal neural stem/progenitor cells. Moreover, TBX1 transcriptionally regulates genes linked to post-embryonic neurogenesis and neurodevelopmental disorders associated with other rare gene variants. Our data indicate a precise time window and cell type through which the social dimension is altered by a gene encoded in a rare CNV and provide a potential common mechanistic basis for a group of neurodevelopmental disorders.

**One-Sentence Summary:** *Tbx1*, a gene affecting neonatal stem cell proliferation, influences peripubertal social behavior.

## Main Text

Identifying the developmental origin of neurodevelopmental disorders and the cell types in which gene variants cause such disorders is essential to develop mechanism-based precision medicine in psychiatry. Rare copy number variants (CNVs) provide a reliable entry point to delve into the pathophysiology of high-penetrance genetic risk factors, linking genes to cellular phenotypes and behavioral consequences. However, CNVs pose conceptual and technical challenges, as each CNV is associated with multiple mental illnesses. A technical challenge is that any CNV might encode many functionally diverse genes. In theory, to determine how CNV-encoded single genes affect neurodevelopmental disorders, isolated dose manipulation of each CNV-encoded single gene (or a set of such genes) and identification of the resulting phenotypes are required. However, such experimental manipulation is not feasible in humans.

As an alternative strategy, CNV-encoded driver genes can be inferred from the protein-truncating variants of each single gene in disease cases. Recent large-scale exome-sequencing studies have identified ultra-rare variants of single genes encoded in CNVs among individuals with schizophrenia and autism spectrum disorder (ASD) (*1, 2*). However, the failure to identify such variants does not necessarily prove their absence. In fact, small-scale exome-sequencing studies have not identified ultra-rare variants that larger scale analyses have. Moreover, ultra-rare variants may also exist in introns and promoter/enhancers of CNV-encoded single genes (*2, 3*). Further, it is possible that such variants simply do not exist in nature but a one-copy deletion of such single genes within a CNV produces certain phenotypes.

Hemizygous deletion at human chromosome 22q11.2 is associated with elevated rates of schizophrenia and diverse neurodevelopmental disorders, including ASD, intellectual disability, and attention-deficit hyperactivity disorder (*4*). Ultra-rare cases of protein-truncating variants of *TBX1* have also been reported in some families, and some of those carriers have been diagnosed with ASD (*5, 6*). However, these individuals also carry variants of genes outside 22q11.2, precluding the establishment of a causative role for this gene variant (*5*). A recent large-scale exome-sequencing study of schizophrenia cases identified a protein-truncating variant of *TBX1* (*1*). However, power limitations of these human studies render the contribution of *TBX1* to mental illnesses or their dimensions uncertain.

The causal relation between CNVs and phenotypes can also be explored along dimensions crossing clinical boundaries (*7*). Many dimensions of social and cognitive domains common to ASD and schizophrenia are negatively impacted by CNVs (*8, 9*). While animal models do not recapitulate clinically defined mental disorders, a dimensional analysis provides an opportunity to translate human phenotypes to those in experimental animals. Complementing the limitations of human studies, reports on genetic mouse models for single CNV-encoded genes have suggested that some 22q11.2-encoded genes contribute to distinct dimensional aspects of mental illnesses (*10*). Constitutive heterozygosity of *Tbx1* in a homogeneous genetic background impairs social communication and interaction in mice (*11- 13*). We capitalized on these observations to further identify the precise developmental time window, critical cell type, and functions associated with *Tbx1* in mouse and cell models.

## RESULTS

### Critical developmental time points of Tbx1 for hippocampal neurogenesis

Human *TBX1* and its murine homolog *Tbx1* are expressed in the brains of humans (*14*) and mice (*15*), respectively. TBX1 protein levels in the whole mouse brain and hippocampus precipitously declined from the embryonic period to adulthood (**Fig. S1A–C**). Constitutive *Tbx1* heterozygosity alters cortical neurogenesis at embryonic day 13.5 (E13.5) in some, but not all cortical regions (*16, 17*). In the hippocampus of the adult mouse, TBX1 protein is more enriched in the granule cell layer of the hippocampus than in the cortex (*11*). We examined the rate of embryonic neurogenesis in the hippocampus at E18, as embryonic neurogenesis in the hippocampal granule cell layer peaks around E17–E19 in rodents (*18*). BrdU injected at E18 labeled indistinguishable numbers of cells in various hippocampal regions in *Tbx1*^+/+^ and *Tbx1*^+/-^ mice (**Fig. S2**), indicating that *Tbx1* heterozygosity had no detectable impact on embryonic neurogenesis in the hippocampus.

The granule cell layer of the hippocampus undergoes protracted neurogenesis, up to postnatal day 18 (P18), and a small fraction of neurogenesis persists in the subgranular zone toward adulthood (*18*). However, the number of BrdU-positive cells in the hippocampus was statistically indistinguishable between *Tbx1*^+/+^ and *Tbx1*^+/-^ mice at P35 (**Fig. S3**).

Several interpretative issues are associated with BrdU (*19, 20*). A single dose of BrdU labels the DNA synthesis of S-phase cells at the time of injection, but it is not a marker of dividing cells *per se*. While a higher dose or repeated injections of BrdU would label more cells, it could additionally label cell death or repair. Moreover, the number of BrdU-labeled cells could be confounded by altered lengths of the S-phase and total cell cycle. Thus, to reveal the actual neural stem/progenitor cells during the postnatal period, we developed Tg(Nes-EGFP)33Enik/J;*Tbx1*^+/-^ mice (see Supplementary Information) and counted the numbers of GFP-positive neural stem/progenitor cells in the subgranular zone at P35. There were significantly fewer GFP-positive cells in the subgranular zone of the dorsal hippocampus in Tg(Nes-EGFP)33Enik/J;*Tbx1*^+/-^ mice than in Tg(Nes-EGFP)33Enik/J;*Tbx1*^+/+^ mice (**Fig. 1**). Thus, *Tbx1* heterozygosity reduces the number of stem/progenitor cells in the postnatal hippocampus.

**Fig. 1.**
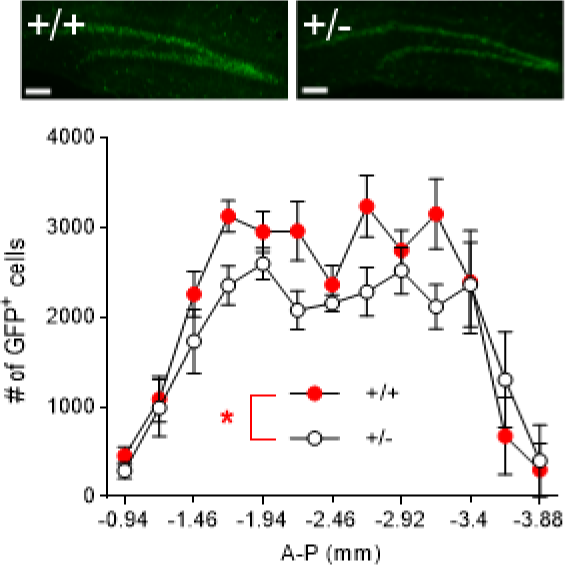
Effect of constitutive *Tbx1* heterozygosity on adult neural stem/progenitor cells in the mouse hippocampus. (**A**) Representative images of GFP-positive cells in the subgranular zone of the hippocampus of Tg(Nes-EGFP)33Enik/J;*Tbx1^+/+^*and Tg(Nes-EGFP)33Enik/J;*Tbx1^+/-^* mice at P35. The scale bar indicates 100 µm. (**B**) More GFP-positive cells were found in the subgranular zone of the hippocampus of Tg(Nes-EGFP)33Enik/J;*Tbx1^+/+^* mice than in Tg(Nes-EGFP)33Enik/J;*Tbx1^+/-^* mice (genotype, F(1,9) = 9.209, *P* = 0.014; A-P position, F(12,108) = 16.92, *P* < 0.0001; genotype × A-P position, F(12,108) = 1.119, *P* = 0.353). For clarity, data are presented as means ± SEM but were analyzed using a generalized linear regression model. Tg(Nes-EGFP)33Enik/J;*Tbx1^+/+^* mice, *N* = 6; Tg(Nes-EGFP)33Enik/J;*Tbx1^+/-^* mice, *N* = 5. In total, 13 sections were counted per animal.

### Cellular consequences of Tbx1 deficiency in neonatal neural stem/progenitor cells

We further explored the precise manner through which *Tbx1* heterozygosity impacts the stem/progenitor cell population. To determine the cell-autonomous effect of *Tbx1* deficiency on the cell cycle of stem/progenitor cells, we cultured and isolated stem/progenitor cells derived from the P0 hippocampus of C57BL/6J pups. This developmental time point was chosen because the TBX1 protein level is higher in the hippocampus earlier than later in the post-embryonic period (see **Fig. S1C**), and constitutive *Tbx1* heterozygous mice show defective social communication as early as P7–8 (*11-13*). Cells were made to simultaneously enter the cell cycle simultaneously using the double-thymidine block method (*21*). TBX1 protein levels peaked during the G1/S phase shortly before the levels of cyclin B1—a marker of the G2/M phase of the cell cycle—peaked (**Fig. 2A**).

**Fig. 2.**
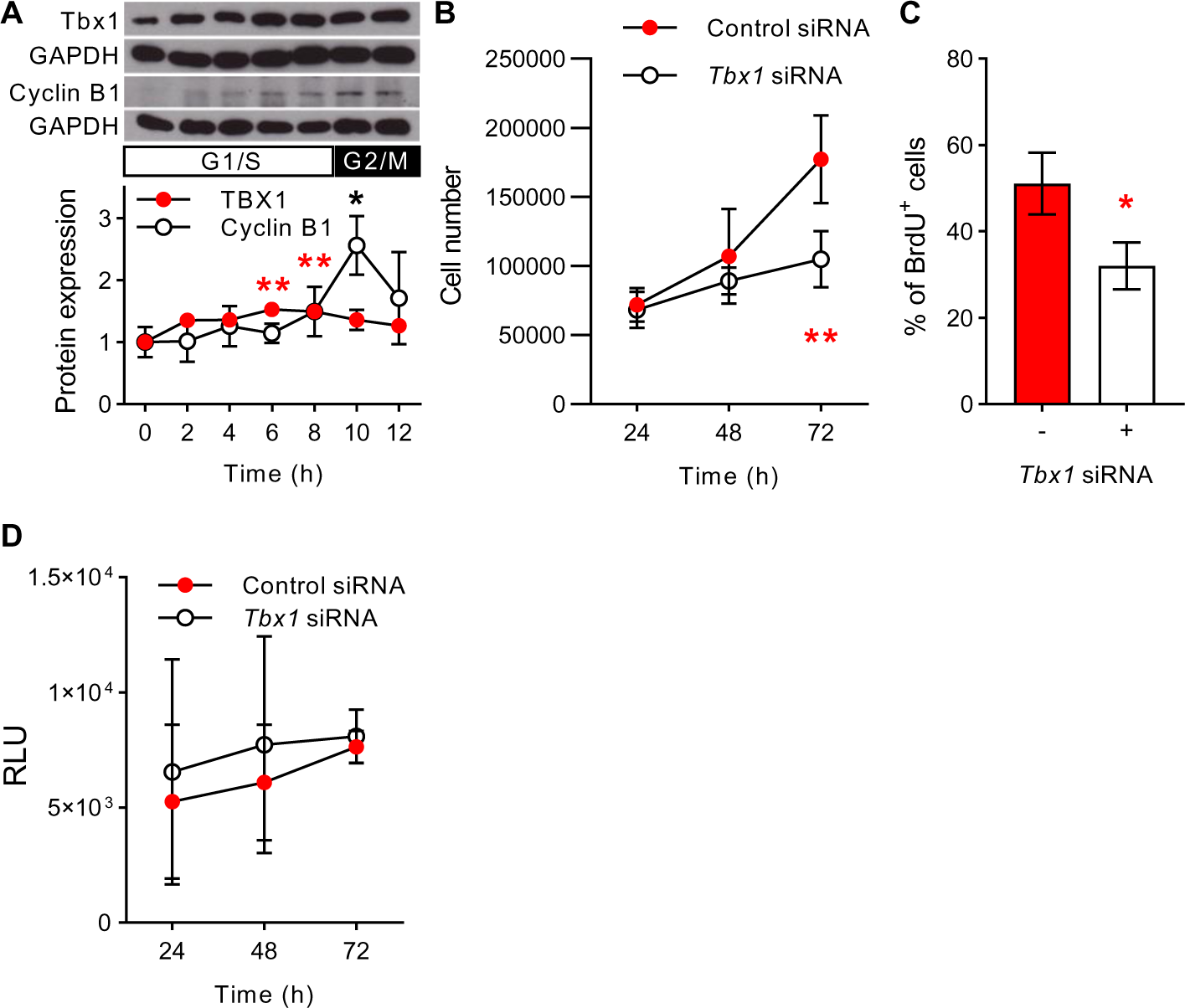
Effect of *Tbx1* deficiency on neonatal neural stem/progenitor cells. (**A**) TBX1 significantly increased at 6 and 8 h, compared to 0 h (\*\**P* < 0.05). Cyclin B1 expression levels peaked at 10 h, compared to 0 h (\**P* < 0.05). Protein expression levels were calculated by dividing the intensity of TBX1 or cyclin B1 signals by that of GAPDH. TBX1; *N* = 6 per time point, except for *N* = 4 at 12 h. Cyclin B1; *N* = 6 per time point, except for *N* = 5 at 6 h and *N* = 4 at 12 h). (**B**) *Tbx1* siRNA blunted neonatal stem/progenitor cell proliferation at 72 h (\*\**P* = 0.001). *N* = 4, 5, and 7 at 24, 48, and 72 h, respectively, for Control siRNA and *Tbx1* siRNA. (**C**) *Tbx1* siRNA significantly reduced BrdU-positive cell number (\**P* = 0.04; Control siRNA *N* = 9, *Tbx1* siRNA *N* = 9). (**D**) *Tbx1* siRNA did not alter the apoptosis rate of neonatal stem/progenitor cells, as determined by the RealTime-Glo Annexin V assay (*P* > 0.05 for each time point). RLU, relative light unit. Control siRNA *N* = 9 and *Tbx1* siRNA *N* = 9 for each time point.

When we reduced TBX1 expression level using siRNA *in vitro* (**Fig. S1B**) and examined the rate of proliferation of neonatal neural stem/progenitor cells under proliferating conditions, *Tbx1* siRNA significantly reduced the cell proliferation rate (**Fig. 2B**). As this is a selective population of neonatal neural stem/progenitor cells under a maximally proliferative condition *in vitro*, the detection sensitivity of BrdU is expected to be higher than that of *in vivo* samples. *Tbx1* siRNA reduced the number of BrdU-positive cells (**Fig. 2C; Fig. S4**). Moreover, the reduced proliferation rate was not due to apoptosis (**Fig. 2D**). Thus, TBX1 deficiency cell-autonomously reduces the proliferation rate of neonatal hippocampal neural stem/progenitor cells.

### Behavioral consequences of Tbx1 deficiency in neonatal neural stem/progenitor cells

Constitutive *Tbx1* heterozygous pups exhibit defective neonatal social communication as early as P7–8 (*11, 13*), and *Tbx1* deficiency blunts the proliferation of neural stem/progenitor cells derived from the P0 hippocampus (see **Fig. 2B**). Therefore, we investigated whether *Tbx1* insufficiency during the neonatal period is critical for the normal development of peripubertal social behavior. To initiate *Tbx1* heterozygosity in neural stem/progenitor cells at specific post-embryonic periods, we developed nestinCreERT;*Tbx1*^flox/+^ mice (c*Tbx1*^+/-^ mice) (see Supplementary Information). *Tbx1* heterozygosity was initiated by tamoxifen at P1–5 and mice were behaviorally tested 1 month later (**Fig. 3A**); it takes at least 1 month for new neurons to be incorporated into the hippocampal circuitry in rodents (*22*). Recombination was found to be selectively induced in newly generated neurons in brain regions with known post-embryonic neurogenesis in response to tamoxifen (**Fig. S6**). As vehicle-treated nestinCreERT;*Tbx1*^flox^ mice and tamoxifen-treated wild-type;*Tbx1*^flox^ mice did not differ in social behavior, they were combined as a single control group (**Fig. S5A–E**). c*Tbx1*^+/-^ mice showed a social approach deficit (**Fig. 3B**), with normal levels of approach behavior toward a novel object (**Fig. 3C**), anxiety-related traits (**Fig. 3D**), and motor behavior (**Fig. 3E**). The predominant form of social approach was sniffing; therefore, we additionally investigated whether the social approach deficit in c*Tbx1*^+/-^ mice was due to their abnormal reaction to social olfactory cues. Control and c*Tbx1*^+/-^ mice were indistinguishable in their responses and habituation to various non-social and social odorants (**Fig. S7AB**). These data show that the reduced levels of social approach in c*Tbx1*^+/-^ mice in the peripubertal period (i.e., 1 month of age) are not due to the generalized deficits in olfactory responses to social cues, novel object approach, anxiety, or motor capacity.

**Fig. 3.**
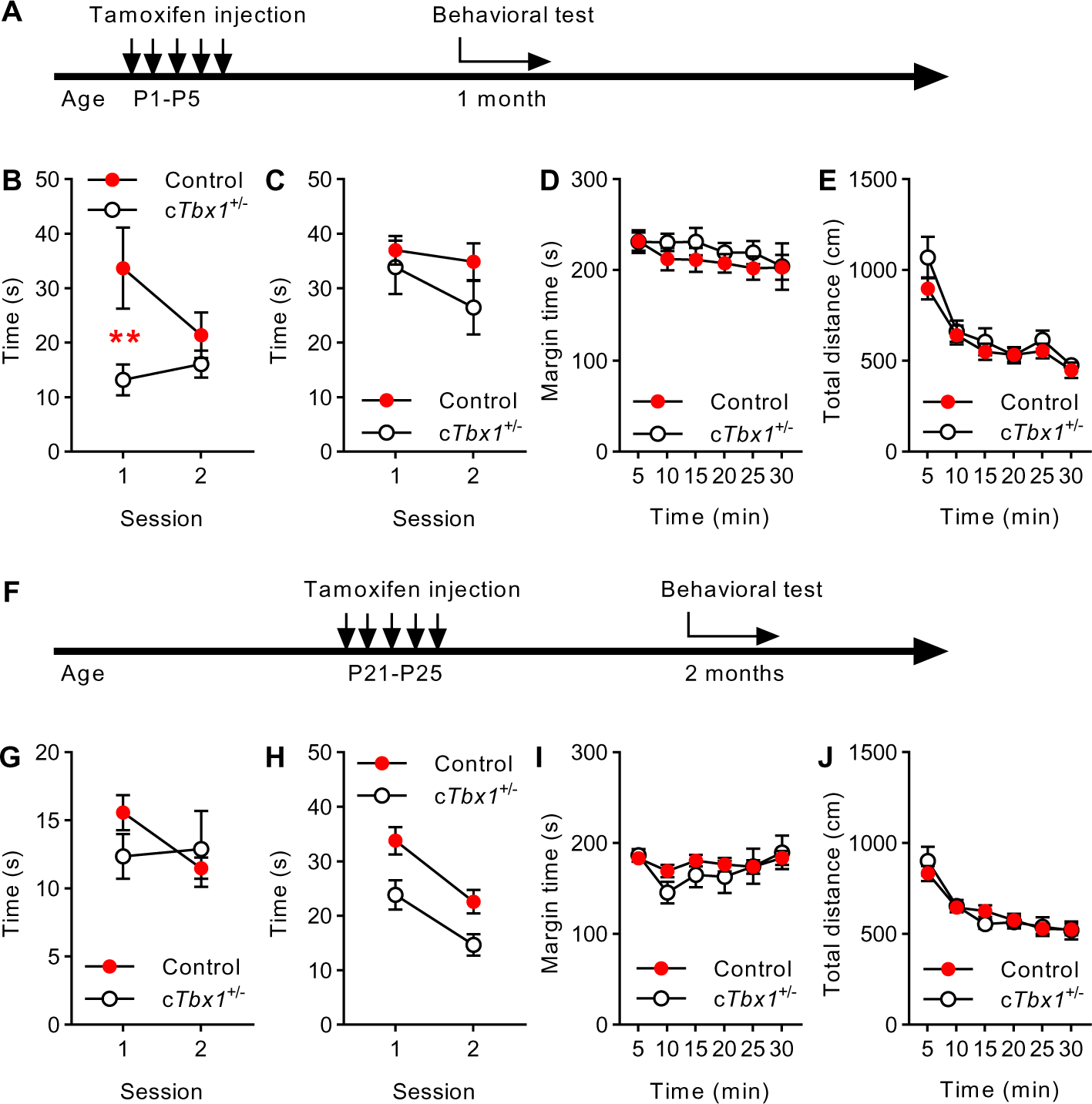
Behavioral effect of *Tbx1* deletion initiated during neonatal and postnatal periods. Tamoxifen (0 and 83.5 mg/kg body weight, i.p.) was given to lactating mothers of nestinCreERT;*Tbx1^flox/+^* pups from P1 to P5. Behavioral assays started 1 month later (**A**) for social approach (**B**) (Control, *N* = 19; c*Tbx1^+/-^*, *N* = 12), novel object approach (**C**) (Control, *N* = 23; c*Tbx1^+/-^*, *N* = 12), thigmotaxis (**D**) (Control, *N* = 14; c*Tbx1^+/-^*, *N* = 6), and motor activity (**E**) (Control, *N* = 14; c*Tbx1^+/-^*, *N* = 6). c*Tbx1^+/-^* mice showed lower levels of social approach at session 1 than control mice did (*P* = 0.007). The groups did not differ in any other tested behavior (*P* > 0.05). Another set of mice were given tamoxifen (0 and 180 mg/kg body weight, i.p.) from P21 to P25 and were tested 1 month later (**F**) for social approach (**G**) (Control, *N* = 35; c*Tbx1^+/-^*, *N* = 11), novel object approach (**H**) (Control, *N* = 33; c*Tbx1^+/-^*, *N* = 12), thigmotaxis (**I**) Control, *N* = 40; c*Tbx1^+/-^*, *N* = 12), and motor activity (**J**) (Control, *N* = 40; c*Tbx1^+/-^*, *N* = 12). c*Tbx1^+/-^* mice showed lower levels of novel object approach equally at sessions 1 and 2 compared to control mice (genotype, F(1,43) = 5.151, *P* = 0.028; genotype × session, F(1,43) = 0.635, *P* = 0.43). The groups did not differ in any other tested behavior (*P* > 0.05).

To further examine the specificity of the neonatal period, we initiated tamoxifen treatment at P21–25 in a different group of mice and tested them 1 month later (i.e., at 2 months of age) (**Fig. 3F**); mice of various control genotypes did not differ, and they were combined as a single control group (**Fig. S5F–J**). These c*Tbx1*^+/-^ mice showed no deficits in social approach, an anxiety-related trait, or motor capacity in an open field (**Fig. 3G, I, and J**), although they showed impaired ability to approach a novel non-social object (**Fig. 3H**). That tamoxifen treatment at P21–25 affected novel object approach indicates that this treatment *per se* was effective, ruling out the possibility that the tamoxifen dose was insufficient to impact adult neurogenesis and behavior.

The ablation of postnatal/adult neural stem/progenitor cells at P28 or later has no effect on the subsequent social approach/interaction *per se* in mice (*23-25*). Our observations are consistent with these findings in that *Tbx1* deletion in stem/progenitor cells at P21–25 had no detectable effect on later social approach/interaction in male mice, but further extend the previous studies by showing that neonatal neurogenesis is more critical for the normal development of social behavior than postnatal neurogenesis and Tbx1 functionally contributes to the development of peripubertal social behavior via its actions during the neonatal period.

### Molecular network of *TBX1* in neonatal neural stem/progenitor cells

Having identified neonatal neural stem/progenitor cells as a cellular substrate through which *Tbx1* deficiency causes social approach deficits later, we next examined the molecular targets of this transcription factor using chromatin immunoprecipitation (ChIP)-seq. We determined TBX1-binding sites under a proliferating condition in neonatal stem/progenitor cells derived from the hippocampus of P0 C57BL/6J mice. We ran two sets of ChIP-seq independently using different protocols. TBX1-binding sites were associated mostly with either two genes (65%) or one gene (30%), and in some instances at sites with no nearby genes (5%) (**Fig. S8A**). This analysis collectively identified 1,257 genes and mRNA variants (**Table S1-1**), comprising 1,039 genes at and near the TBX1-binding sites. The distances between the transcription start site and TBX1 binding sites were most frequently found at 50–500 kb in both directions, followed by 5-50 kb, more than 500 kb, and less than 5 kb in this order (**Fig. S8B**). The majority of TBX1-binding sites were intergenic regions (73.42%), followed by introns (12.12%), upstream regions (10.55%), promoters (2.96%), and exons (0.95%) (**Fig. S9**).

Biological annotations showed the enrichment of G-protein–coupled receptor signaling pathways, epigenetic gene regulation, and chromatin silencing among biological processes (**Fig. S10-1**); apical junction complex and cell–cell junctions among cellular components (**Fig. S10-2**); and transcription factor activity, G-protein–coupled activity, structural molecule activity, and transmembrane receptor activity among the molecular functions (**Fig. S10-3**).

Next, we explored databases of genes relevant to adult hippocampal neurogenesis (MANGO v3.2). It should be cautioned that in the absence of a gene database for neonatal neurogenesis, the MANGO database was used as a proxy. Nonetheless, this search identified 30 TBX1-bound genes that were implicated in adult neurogenesis (**Fig. 4A yellow circle**; **Table S1-2**), 22 of which form molecular networks with TBX1 (**Fig. 4B**). We next asked whether our pool of TBX1-bound genes is involved in ASD (**Table S1-3**) and developmental brain disorders (**Table S1-4)**. This search identified 87 genes (**Fig 4A blue circle**). Tbx1 potentially interacts with 61 such genes (**Fig. S11**), among which 12 genes were also listed in the MANGO database, i.e., *Cacna1d*, *Cnr1*, *Dcc*, *Dcx*, *Disc1*, *Dmd*, *Dyrk1a*, *Foxg1*, *Gria1*, *Pten*, *Tbr1*, and *Tdo2* (**Fig. 4A**), rare variants of which are associated with neurodevelopmental disorders, including ASD. Moreover, Tbx1 was found to bind to loci near genes implicated in cell cycle (**Table S1-5**), GABAergic and inhibitory, and glycinergic synapses (**Table S1-6, S1-7, and S1-8**) and glutamatergic and excitatory synapses (**Table S1-9 and S1-10**).

**Fig. 4.**
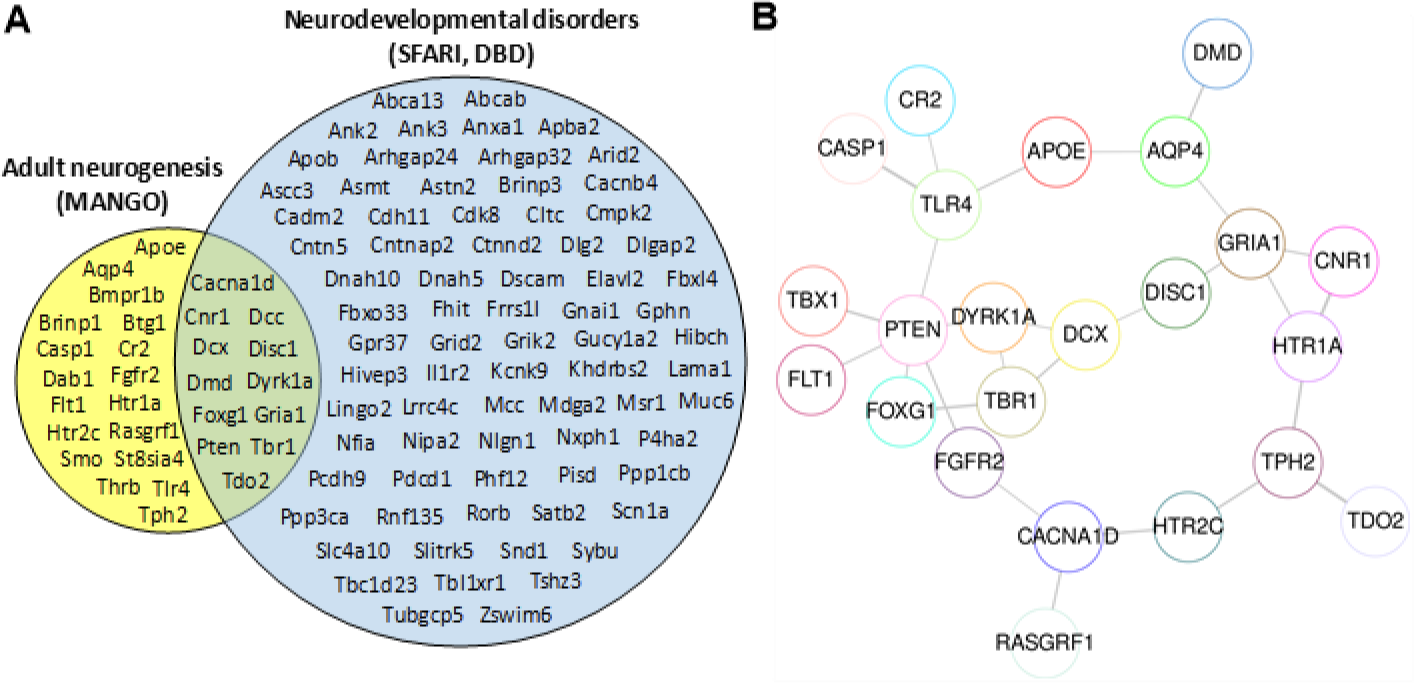
Genes bound by TBX1 that are implicated in adult neurogenesis and neurodevelopmental disorders. (**A**) The following gene databases were used: (http://mango.adult-neurogenesis.de/documents/annotations?show=20&expression=true; MANGO, left yellow circle). Autism and other neurodevelopmental disorders (SFARI, https://gene.sfari.org; DBD, https://dbd.geisingeradmi.org/) (Right, pale blue circle). (**B)** Potential interactions among the target genes of TBX1 are shown, based on Search Tool for Retrieval of Interacting Genes/Proteins (STRING v11) analysis of MANGO-listed genes (see yellow circle in **A**) (*41*).

Molecular networks are based on lenient inclusive criteria; the Search Tool for Retrieval of Interacting Genes/Proteins (STRING) database contains any association among genes or molecules as long as they are mentioned in the same articles (*26*). While such networks are useful for their heuristic value, they may not represent real, functional associations. For example, while a TBX1–PTEN network was identified using the databases of adult neurogenesis (**Fig. 4B**) and neurodevelopmental disorders (**Fig. S11**), this association was not based on any functional validation. Thus, we performed a series of validations. Our ChIP-seq analysis identified the actual binding sites of TBX1 near PTEN (**Table S1-1**). TBX1 was colocalized with PTEN in neonatal neural stem/progenitor cells derived from the P0 hippocampus of C57BL/6J pups (**Fig. 5A**, left) and in a subpopulation of PTEN-positive cells in the hippocampal granular zone of C57BL/6J mice (**Fig. 5A**, right). The *Pten* promoter was enriched in DNA fragments precipitated by a TBX1 antibody, as determined by ChIP-PCR (**Fig. 5B**). *Tbx1* siRNA significantly reduced *Pten* promoter activity (**Fig. 5C**) and PTEN protein levels (**Fig. 5D**). TBX1 protein was found to be bound 315 bp upstream and 84,805 downstream of the *Pten* transcription start site (**Fig. S12A–B**). The former site is within a reasonable enhancer location distance (*27*). However, the binding locations of transcription factors revealed by ChIP-seq are not reliable in terms of the precise distances from the target genes owing to the size of fragments generated by sonication (*28*). Moreover, the abundance of binding may not necessarily indicate functionally critical loci. Thus, we constructed various promoter sequences of *Pten* with deletions and mutations at different loci (**Fig. 5E**, left) and examined their transcriptional activity. While the *Pten* promoter is constitutively active, its transcriptional activity doubles during proliferation with epidermal growth factor (EGF) (see **Fig. S13**). The presence of the full-length *Pten* promoter was essential for luciferase activity (**Fig. S14**). The transcriptional activation of the *Pten* promoter under this condition was reduced by a deletion or mutation placed at the proximal 35 bp segment (**Fig. 5E**, right), which contains a recognition sequence resembling the T-box consensus sequence (*29*). Our observations that *Tbx1* deficiency results in a blunted increase in the number of neonatal hippocampal neural stem/progenitor cells (see **Fig. 2B**) and reduction in the transcriptional activation and expression levels of PTEN (see **Fig. 5C–D**) are consistent with the eventual depletion of stem/progenitor cells in the adult hippocampal granule cell layer following *Pten* deletion *in vivo* (*30*). Molecular networks identified by STRING uncovered *Flt1*, *ApoE*, and *Tdo2* as potential indirect interacting molecules that are also implicated in adult neurogenesis (**Fig. 4AB**). While TBX1-binding sites are outside a 2 kb promoter region (**Table S1-1**), *Flt1* was identified in four separate clones of ChIP-seq and thus included. ApoE has not been implicated in neurodevelopmental disorders, but its indirect link with TBX1 via PTEN is a possibility (see **Fig. 4B**). Tryptophan 2,3-dioxygenase (TDO2) is the rate-limiting enzyme in the catabolism of tryptophan in the kynurenine pathway, the downregulation of which has been implicated in ASD (*31*). We additionally tested *Btg1*, as it was identified in MANGO (**Table S1-2**) but not in databases of neurodevelopmental disorders (see **Table S1-3; S1-4**). We reduced *Tbx1* mRNA expression levels in the culture of neonatal neural stem/progenitor cells using *Tbx1* siRNA. This analysis showed that *Tbx1* siRNA reduced *Tbx1* mRNA by half (**Fig. S15A**) and reduced *Tdo2* mRNA levels but did not affect levels of *Flt1*, *ApoE*, or *Btg1* mRNA (**Fig. S15B–E**). *Tdo2* mRNA is also highly enriched in the hippocampal granule cell layer in the mouse brain (*32*) and is critical for the proliferation of adult neural stem/progenitor cells (*33*).

**Fig. 5.**
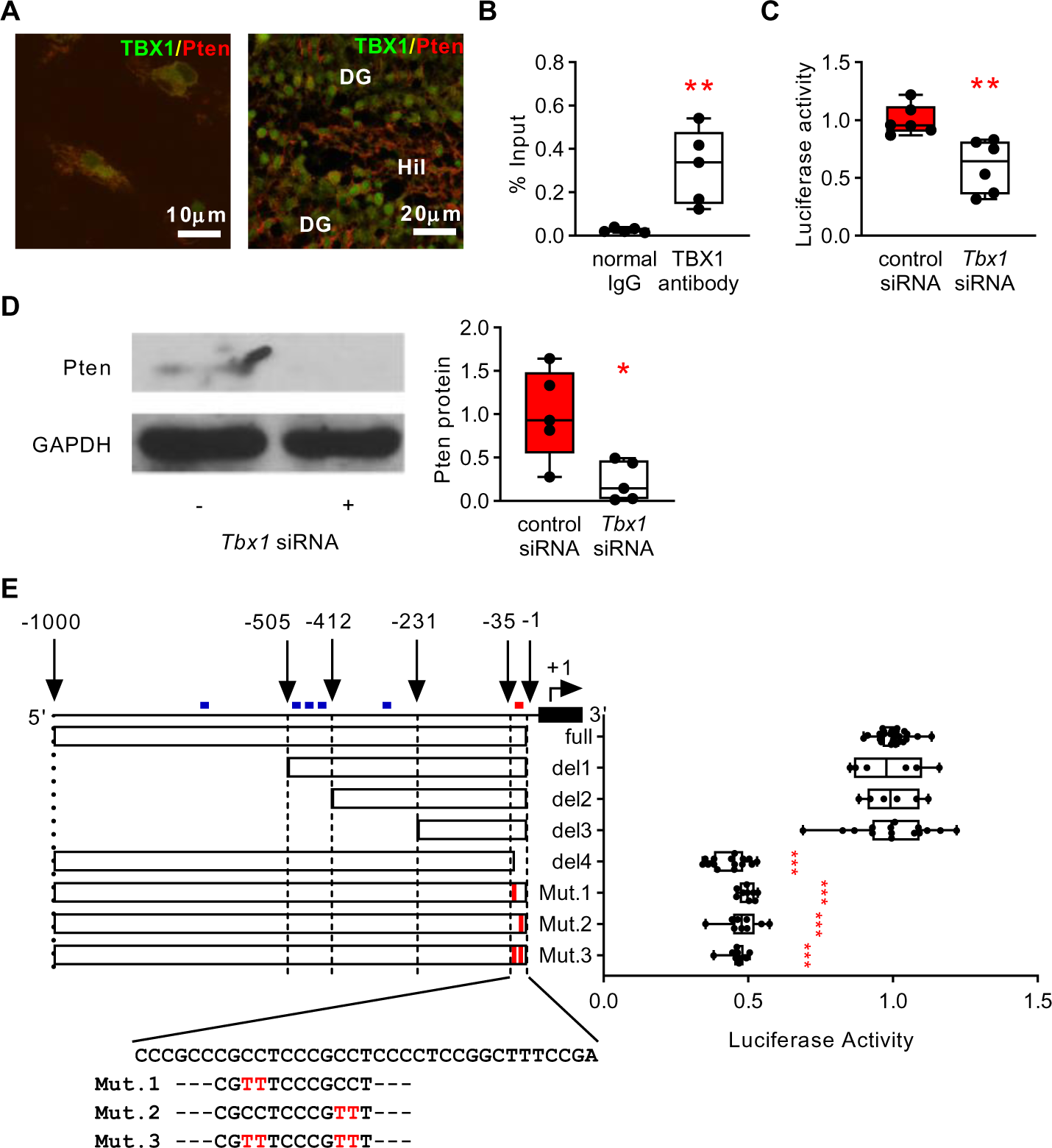
Transcriptional regulation of *Pten* by TBX1 in neonatal stem/progenitor cells. (**A**) TBX1 expression in PTEN-positive stem/progenitor cells *in vitro* (left) and granule cells in the mouse hippocampus (right). (**B**) ChIP-PCR. TBX1 binds to the *Pten* promoter. Input, pre-immunoprecipitated diluted solution was the control (*P* = 0.008; *N* = 5 per group). %input = 2% × 2(C[T] 2%Input Sample − C[T] IP Sample). (**C**) *Tbx1* siRNA significantly reduced transcriptional activity of the 1 kb *Pten* promoter (*P* = 0.002; *N* = 6 per group) (see **Fig. S1B** for *Tbx1* siRNA efficacy). (**D**) *Tbx1* siRNA significantly reduced PTEN protein in P0 neural progenitor cells (*P* = 0.0016; *N* = 5 per group). (**E**) Deletion and mutation positions within the 1 kb *Pten* promoter (left); their effects on luciferase activity (right). Deletion and mutation of the proximal 35 bp reduced *Pten* promoter activity (\*\*\**P* < 0.001; Mann–Whitney test). T-box-binding consensus sequence (blue dots): GTGTTATGACACGTGCAAGTGTGAGTGCGA (*42*); T-box–binding-like sequence (red dot): CCGCCTCCCGCCT (*29*). Full, *N* = 27; Del.1, *N* = 6; Del. 2, *N* = 6; Del. 3, *N* = 15; Del. 4, *N* = 15; Mut.1, *N* = 9; Mut. 2, *N* = 9; Mut. 3, *N* = 9.

TBX1 regulates the proliferation of neonatal neural stem/progenitor cells (see **Fig. 2B**), and *Pten* deletion leads to a transitory expansion of hippocampal adult stem/progenitor cells and other stem cells, followed by the exhaustion of the precursor pool, presumably through an imbalanced increase in the extent of G0 to G1 transition (*34, 35*). Therefore, we further explored the potential target genes implicated in the G0/G1 phase among ChIPseq-identified TBX1-bound genes, which revealed *Ecrg4* and *Rnf112* (**Table S1-1; S1-5**). TBX1-binding sites were found near *Ecrg4* and *Rnf112* (**Fig. S17**–**S18; Table S1-1 and S1-5**). *Tbx1* siRNA significantly reduced the expression of *Ecrg4* and *Rnf112* mRNA (**Fig. S15F–G**). No functional roles of these genes in neonatal neural stem/progenitor cells in the hippocampus are known. However, ECRG4 suppresses the proliferation of stem cells in the mouse dentate gyrus (*36*). Our data identify potential molecular targets whereby TBX1 regulates the cell cycle in hippocampal neonatal neural stem/progenitor cells.

## DISCUSSION

We established that neonatal *Tbx1* deficiency impairs peripubertal social behavior and impairs the proliferation of neonatal neural stem/progenitor cells. Therefore, our study provides a potential cellular basis for a social dimension of neurodevelopmental disorders linked to *TBX1* variants and a critical developmental time point at which TBX1 determines the level of a subsequent social dimension. This gene transcriptionally acts on 87 genes that are independently associated with neurodevelopmental disorders; 12 of these genes are also known to be involved in adult neurogenesis. Interestingly, 75 of the 87 genes associated with neurodevelopmental disorders have not been implicated in adult or neonatal neurogenesis. Moreover, 18 TBX1-bound genes that are listed in the adult neurogenesis database have not been known to be associated with neurodevelopmental disorders. This pool of genes provides a means to identify novel genes contributing to the social dimension of neurodevelopmental disorders via neonatal neurogenesis. Thus, our data link *TBX1*, which has its own ultra-rare gene variants and is also encoded in a rare CNV, to a set of other genes, rare variants of which are associated with neurodevelopmental disorders.

How CNV-encoded genes and their biological processes contribute to various psychiatric disorders is still poorly understood. The identification of driver genes within CNVs in humans is limited due to the very rare nature of variants of their encoded genes. Recent large-scale exome sequencing studies have identified more ultra-rare variants (*1, 37*), but these data do not necessarily rule out the possibility that dose alterations--not their own variants—of CNV-encoded genes contribute to phenotypes. Moreover, while embryonic neurogenesis, synaptic functions, and activity-regulated cytoskeleton-associated proteins have emerged as common, robust candidate processes affecting mental illnesses, such analyses are not devoid of technical and interpretative limitations (*38*). Literature-curated databases do not provide genes and biological processes whose functions have not been explored or are not abundantly or commonly represented.

Our approach, starting from a CNV-encoded gene, represents a complementary strategy to circumvent the limitations associated with human studies and to identify driver genes, and their cell functions and dimensional behavioral phenotypes. We capitalized on TBX1, a 22q11.2 CNV-encoded gene, of which several cases of ultra-rare variants exist with variable neurodevelopmental disorders (*1, 5, 6, 39, 40*). The causative roles of *TBX1* in phenotypes have not been established due to their lack of statistical power (*1*) and the presence of other variants among carriers (*5*) in humans. Given that the expression of TBX1 protein is enriched in the sites of post-embryonic neurogenesis (*11*), we focused on the function of this gene in post-embryonic stem cells in mice. This approach provided a theoretical basis for grouping dimensional aspects of neurodevelopmental disorders, based on a shared mechanistic basis, a prerequisite for designing mechanism-based precision medicine to ultimately treat specific dimensions of such disorders in humans (*7*).

## Acknowledgments

We thank Dr. Bernice Morrow for providing the original breeders of *Tbx1*^+/-^ and *Tbx1*^flox/+^ mice.

## Author Contributions

Conceptualization: TH, NH

Methodology: TH, SB, GK, SA, KT, MN, TVM, GE

Investigation: TH, NH, BS, GK, SA, KT, MN, MBS, PÓ, KY,

Visualization: KY, TY

Funding acquisition: NH, SB

Project administration: NH

Supervision: NH

Writing – original draft: TH, SB, GK, PÓ, MBS, NH

Writing – review & editing: TH, SB, NH

## Data and material availability

All data are deposited at https://datadryad.org/stash/. Mice are available through material transfer agreement.

## Supplementary Materials

### Materials and Methods

#### Experimental Design

The objective of this study was to determine the developmental time point(s), cell types, and target genes of the transcription factor TBX1 required for the normal development of social behavior in mice. The study design included four major components. First, the impact of constitutive *Tbx1* heterozygosity on hippocampal embryonic and postnatal neural stem/progenitor cells was examined in mice. Second, we determined the cell-autonomous effect of *Tbx1* deficiency on the proliferation and apoptosis of neonatal neural stem/progenitor cells *in vitro*. Third, we determined the impact of *Tbx1* heterozygosity initiated at P1–5 and P21–25 on social and other behaviors. Fourth, we identified TBX1- binding sites in the whole genome using ChIP-seq and neonatal stem/progenitor cells of the hippocampus and validated the potential target genes of TBX1.

Details of the primer sequences used for qRT-PCR analysis (Fig. S15) are listed in Table S3. The validation of the antibodies and reagents is provided in Tables S4 and S5, respectively.

#### Mice

The protocols for animal handling and experimentation were approved by the Animal Care and Use Committee of the Albert Einstein College of Medicine, Shiga Medical Center (Approval number 24-2), and the University of Texas Health Science Center at San Antonio (20190084AR), in accordance with the NIH guidelines. Genotypes were determined by PCR analysis of samples obtained from tail tissue. Male littermates were used in this study.

##### C57BL/6J and C57BL/6N mice

We purchased C57BL/6J mice from the Jackson Laboratory (Stock No. 000664, Bar Harbor, ME, USA) and C57BL/6N from SLC (Hamamatsu, Japan).

##### *Tbx1*^+/-^ mouse

This mouse was a constitutive *Tbx1* heterozygous mouse, with more than 10 generations of backcrossing to C57BL/6J mice (*11*). For BrdU labeling of embryonic mice only, we used congenic *Tbx1*^+/-^ mice with more than 10 generations of backcrossing to C57BL/6N.

##### Nestin-GFP;*Tbx1*^+/-^ mouse

We backcrossed Tg(Nes-EGFP) mice (*43*) to C57BL/6J mice for 10 generations and then crossed them with our *Tbx1*^+/-^ mice.

##### c*Tbx1*^+/-^ mouse

C57BL/6-Tg(Nes-cre/ERT2)KEisc/J mice (Nes-cre/ERT2; Cat#016261, Jackson Laboratory) (*44*) and *Tbx1*^flox/+^ mice (*45*) were backcrossed to C57BL/6J for more than 10 generations. In this Nes-cre/ERT2 mouse, Cre-triggered secondary abnormalities do not occur, unlike other Cre mice with the nestin promoter (*44*), and tamoxifen does not cause recombination outside the central nervous system (*46*). C57BL/6-Tg(Nes-cre/ERT2)Keisc/J mice express Cre in adult neural stem/progenitor cells with high specificity and without leakiness. The selectivity of Cre expression in stem/progenitor cells in these mice has been demonstrated as early as the first neonatal week (*47*). Male congenic C57BL/6-Tg(Nes-cre/ERT2)Keisc/J mice or C57BL/6-Tg(Nes-cre/ERT2)Keisc/J;*Tbx1*^flox/+^ mice were crossed with either female congenic *Tbx1*^flox/flox^ or *Tbx1*^flox/+^ mice.

As 3-day (P2–4) administration of tamoxifen to mothers did not achieve full recombination (*47*), we administered tamoxifen for 5 days from P1 to P5. Tamoxifen (#T5648; Sigma, St. Louis, MO, USA) was prepared by vortexing and sonicating 6.7 mg tamoxifen in 100 µL ethanol in a tube wrapped in aluminum foil, followed by the addition of 900 µL sunflower oil and sonication. Tamoxifen at 12.5 µL per gram body weight (0 or 83.5 mg/kg body weight) was intraperitoneally (i.p.) injected into lactating mothers (at the same time of the day) for 5 days; tamoxifen can be delivered to pups through the mother’s milk (*47*). Oral tamoxifen is devoid of adverse effects on glucose tolerance and body composition (*48*). We first compared the control groups with sufficient sample sizes available (tamoxifen wild-type [WT];*Tbx1^flox/+^*, vehicle Nes-cre/ERT2;*Tbx1^flox/+^*, and vehicle Nes-cre/ERT2;*Tbx1^flox/flox^*) to monitor the effects of tamoxifen and nestin-cre/ERT2 on behavior. After no group differences were found, the groups were combined as a single control group (see **Fig. S6A**).

In a separate group of mice, tamoxifen (0 or 180 mg/kg, i.p. per day) was directly injected into Nes-cre/ERT2;Tbx1^flox/+^ from P21 to P25. To monitor the side effects of i.p. injected tamoxifen (48), we compared the following control groups with sufficient sample sizes: tamoxifen-treated wild-type (WT);Tbx1^flox/+^ mice, vehicle-treated Nes-cre/ERT2;Tbx1^flox/+^, vehicle-treated Nes-cre/ERT2;Tbx1^flox/flox^ mice, vehicle-treated WT;Tbx1^flox/+^ mice, vehicle-treated WT;Tbx1^flox/flox^ mice, and tamoxifen-treated WT;Tbx1^flox/flox^ mice. After no group differences were found, the groups were combined as a single control group (see Fig. S6B).

A primary comparison was made between c*Tbx1^+/-^* mice (tamoxifen-treated Nes-cre/ERT2;*Tbx1*^flox/+^) and the combined control.

##### Nes-cre/ERT2 recombination reporter mouse

While the selective recombination in Nes-cre/ERT2 (#016261; Jax, Bar Harbor, ME, USA) in neural stem/progenitor cells during the postnatal and adult periods has been well characterized (*44*, *49*), it has not been after neonatal tamoxifen treatments. We crossed Nes-cre/ERT2 (#016261; Jax, Bar Harbor, ME, USA) with B6.Cg-*Gt(ROSA)26Sor^tm3(CAG-EYFP)Hze^*/J (Ai3(RCL-EYFP, #007903; Jax). Mice were given tamoxifen at P1- to P5 and sacrificed 1 month later. This is the time point when neonatal stem/progenitor cells are expected to be mostly differentiated into mature neurons within the hippocampal granular cell layer. EYFP protein was stained with a GFP antibody (see <u>Immunohistochemistry</u> below). GFP signals were selectively found in the subgranular zone of the hippocampus and the subventricular zone; GFP signals were partially and mostly colocalized with DCX- and NeuN-positive cells, respectively (see **Fig. S5**, DXC/GFP, NeuN/GFP). This pattern is consistent with what is expected to be recombination confined to post-embryonic neural stem/progenitor cells (*44*, *49*). Other brain regions did now show any detectable or consistent GFP signal.

#### Western blotting

To compare TBX1 protein expression levels at different ages and brain regions, brain samples were used for western blotting. Whole brain (10 µg) or hippocampal samples (10 µg) were obtained from male C57BL/6J mice at embryonic day 14, embryonic day 18, postnatal day 7, postnatal day 15, postnatal week 5 (1-month old) and postnatal week 8 (2-month old). Cultured hippocampal stem/progenitor cells (2.5 µg) were derived from P0 C57Bl/6J pups and collected for western blotting 72 h after adding *Tbx1* siRNA.

We used a rabbit monoclonal antibody against TBX1 (1:500, ab109313; Abcam, Cambridge, MA, USA), mouse monoclonal antibody against PTEN (1:100, sc-7974; Santa Cruz), mouse monoclonal antibody against cyclin B1 (1:100; Santa Cruz), mouse monoclonal antibody against GAPDH (1: 500, sc-32233; Santa Cruz), goat anti-rabbit horseradish peroxidase-conjugated IgG secondary antibody (1:2,000, #31460; Pierce, Waltham, MA, USA), and goat anti-mouse horseradish peroxidase-conjugated IgG secondary antibody (1:5,000, #31430; Pierce). The specific bands were visualized using SuperSignal West (#34080; Thermo Fisher). The levels of TBX1 and PTEN were normalized to those of GAPDH. The specific bands were visualized using SuperSignal West. The levels of TBX1 were normalized to those of GAPDH.

#### Cell culture

We followed the standard procedure for culturing neural progenitor cells (*11*, *50*). Briefly, hippocampal tissues were obtained from C57BL/6J pups at postnatal day 0 (P0). Cells were maintained in Dulbecco’s modified Eagle’s medium (DMEM)/F-12 (#11330057; Invitrogen, Waltham, MA, USA) containing N2 (#17502-048; Invitrogen), B27 (#17504-044; Invitrogen), 1% penicillin/streptomycin (#15140122; Invitrogen), and 20 ng/mL epidermal growth factor (EGF) (#236-EG-200; R&D Systems, Minneapolis, MN, USA). Single-cell suspensions of neurospheres were plated on 35 mm dishes coated with poly-L-ornithine (P4957; Sigma) and fibronectin (#1030-FN-05M; R&D) at a density of 4 × 10^5^ cells/dish for western blotting and cell counting or at a density of 2 × 10^5^ cells/dish for fluorescent immunolabeling.

#### siRNA treatment

Cells (2.5 ×10^5^ cells/well in 6-well plates or 35 mm dishes) were grown for 24 h in culture medium before siRNA treatment. Cells were treated with either control siRNA (40 nM, control siRNA-A, sc-37007; Santa Cruz, Dallas, TX, USA) or *Tbx1* siRNA (40 nM, #sc-38468; Santa Cruz) with siLentFect Lipid Reagent (#1703361; Bio-Rad, Hercules, CA, USA) for 24 h. For cell counting, cells were suspended in 150 µL PBS per dish. Trypan blue solution (10 µL, #15250061; Thermo Fisher, Waltham, MA, USA) was added to 10 µL of cell suspension. Ten microliters of the 20 µL mixed solution was loaded into a hemocytometer, and live cells were counted under a 10× objective lens at 24, 48, and 72 h after siRNA transfection. Three clones were used, and the experiment was performed four times, of which two experiments were performed with cells from the same clone. For western blotting and qRT-PCR, cells were collected and analyzed 24, 48, and 72 h after the addition of siRNA. For the BrdU assay, three to five images per dish were randomly captured for cell counting.

#### BrdU

*Tbx1*^+/-^ males were paired with wild-type female *Tbx1*^+/+^ mice as breeders. Pregnant mice were injected with BrdU (50 mg/kg, i.p., B9285; Sigma) at E18.5 *in vivo* and anesthetized with pentobarbital (60 mg/kg, i.p.). The embryos were sacrificed 2 h later, the time point when robust BrdU labeling is detectable (*51*). The brains were post-fixed with 4% paraformaldehyde for 24 h and placed in glycerol for cryoprotection. We used two litters from different mothers. Brains were coronally cut at 40 µm. BrdU-positive cells were counted for each division of the embryonic hippocampus: the dentate gyrus, fimbrio-dentate junction, and dentate ventricular zone. Sections were stained for BrdU according to our standard DAB staining procedure (*11*), and the BrdU-positive cells centered in each division were counted under a light microscope (Zeiss, Jena, Germany).

Five-week-old male *Tbx1*^+/+^ and *Tbx1*^+/-^ mice were injected with BrdU (50 mg/kg, i.p.), anesthetized with pentobarbital (60 mg/kg, i.p.) 24 h later (*52*), and perfused with 4% paraformaldehyde. The brains were coronally sliced at 40 µm, covering the entire antero-posterior region of the hippocampus. Every 6^th^ section was used for stereological counting. Sections were stained for BrdU according to our standard DAB staining procedure (*11*), and the BrdU-positive cells centered in the subgranular zone of the dentate gyrus were counted under a light microscope (Zeiss). BrdU-positive cells were counted for each anterior-posterior (AP) position (from the bregma, AP, −0.94 to −3.88) and for each division.

Single-cell suspensions of neurospheres, derived from P0 C57BL/6J pups, were plated on coverslips coated with poly-L-ornithine (Sigma) and fibronectin (R&D) at a density of 2 × 10^5^ cells/35 mm dish. Twenty-four hours later, siRNA was applied for 24 h. Forty-seven hours after the siRNA was washed away, cells were treated with BrdU (10 μM) and fixed with 4% paraformaldehyde for analysis 1 h later. Cells were nestin-positive under growth conditions. Three to five images were captured from each dish.

Cells were stained for DAPI and BrdU (rat monoclonal BrdU antibody, OBT0030S; Bio-Rad), together with secondary antibodies conjugated with Alexa-594 (1:500, A-11007; Thermo Fisher). The number of DAPI-positive cells and BrdU-positive cells was counted, and the percentage was calculated by dividing the number of BrdU-positive cells by the total number of cells.

#### GFP signals of nestin-GFP;*Tbx1*^+/-^ mice

Five-week-old male nestin-GFP;*Tbx1*^+/+^ and nestin-GFP;*Tbx1*^+/-^ mice were perfused with 4% paraformaldehyde. The brains were coronally sliced at 40 µm, covering the entire antero-posterior region of the hippocampus. Every 6^th^ section was used for the stereological counting. A clustered band of GFP-positive cells in the subgranular zone of the hippocampus was counted under a fluorescence microscope (Zeiss Axioscope). GFP-positive cells were counted for each anterior-posterior position (from the bregma, AP −0.94 to −3.88).

#### Cell cycle synchronization

We used a double-thymidine block method to synchronize the cell cycle (*21*). Briefly, stem/progenitor cells (4 × 10^5^ cells/dish) derived from the P0 hippocampus of C57BL/6J pups were plated and grown overnight. Thymidine (1 mM) was added to the culture medium, and the cells were incubated for 19 h. The medium was replaced with fresh culture medium, and the cells were incubated for 9 h. Finally, thymidine was added to the medium again, and the cells were incubated for 16 h. The medium was replaced with fresh culture medium without thymidine, and the cells were collected 0, 2, 4, 6, 8, 10, and 12 h after the final replacement. TBX1, GAPDH, and cyclin B1 were quantified by western blotting (see Western blotting; Table S4). The relative expression levels of TBX1 and cyclin B1 protein to those of GAPDH were evaluated at the beginning of the cell cycle.

#### Immunohistochemistry

Forty-micrometer-thick sections were obtained from mouse brains. Free-floating sections were stained with a mouse monoclonal anti-PTEN antibody (1:100, sc-7974; Santa Cruz) and a rabbit polyclonal anti-TBX1 antibody (1:100, ab84730; Abcam) together with a goat anti-mouse IgG conjugated with Alexa-594 (1:200; Invitrogen) and goat anti-rabbit IgG conjugated with Alexa-488 (1:200; Invitrogen). Sections were examined using confocal microscopy (see **Fig. 5A**).

The following antibodies were used to localize cells where Cre-triggered recombination took place (see **Fig. S5**): a purified mouse anti-nestin antibody (1:25, BD611658, BD Transduction Laboratories), mouse anti-NeuN antibody (1:200, cat# MAB377, Millipore), goat polyclonal antibody against DCX (1:25, c- 8066; Santa Cruz, Dallas, TX, USA), rabbit polyclonal antibody against GFP (1:400, A11122; Thermo Fisher Scientific), donkey anti-rat IgG conjugated with Alexa-594 (1:200, A-21209; Thermo Fisher), donkey anti-rabbit IgG conjugated with Alexa-488 (1:200, A21206; Thermo Fisher), donkey anti-mouse IgG conjugated with Alexa-594 (1:200, #A21203; Thermo Fisher), donkey anti-mouse IgG conjugated with Alexa Fluor-405 (1:200, ab175658; Abcam), and donkey anti-goat IgG conjugated with Alexa-594 (A11058, #1608643; Thermo Fisher).

To detect GFP signals in tamoxifen-treated B6.129X1-*Gt(ROSA)26Sor^tm1(EYFP)Cos^*/J (R26R-EYFP, #006148; Jax), we stained GFP with an antibody against GFP (rabbit anti-GFP antibody, 1:400, Cat. No. A11122, Life Technologies, Carlsbad, CA, USA) with biotinylated goat anti-rabbit IgG (1:500, Cat. No. BA-1000, Vector Laboratories, Inc., Burlingame, CA, USA) and avidin–biotin complex, using 3,3′-diaminobenzidine (DAB) (*50*).

#### Apoptosis assay

Single cells (0.5 × 10^4^ cells per well; 96-well plate) and culture medium as control were prepared. After siRNA treatment (24, 48, and 72 h), 100 mL 2× Detection reagent (RealTime-Glo Annexin V Apoptosis Assay kit, #JA1000; Promega, Madison, WI, USA) was added and incubated for 30 min. Luminescence was recorded using a GloMax(R) Navigator Microplate Luminometer (#GM2000; Promega). Two clones were used for the five experiments.

#### Behavioral analysis

We used a battery of behavioral assays to characterize non-aggressive affiliative social approach, non-social novel object approach, thigmotaxis, and motor activity (*11*, *50*, *53*). Mice were group-housed as littermates. Testing was done between 10:00 and 16:00 during the light phase of the day. The order of the behavioral assays was based on the imposed stress levels. First, behaviors that occur in cage-like settings (i.e., social approach) and novel object approach were assessed. In tests of social approach, the order of the tested mice was randomized in terms of genotypes. For thigmotaxis and locomotor activity in an open field, a litter of up to four mice was tested simultaneously in four sets of open fields. The sample sizes are provided in the figure legends of the main manuscript. A separate set of mice were used for evaluating the olfactory responses to social and non-social cues.

##### Social approach

A test subject and an unfamiliar stimulus subject (an age-matched male C57BL/6J mouse) were placed in a new home cage setting with new bedding materials. There were no resident mice in this procedure, and the mice did not exhibit aggressive behavior. The assay consisted of two 5 min sessions with 30 min intervals. The total amount of time mice spent exhibiting affiliative non-aggressive approach was analyzed.

##### Non-social novel object approach

This test was used to evaluate the general tendency of a mouse to approach a novel object and habituate to the object following the first encounter. Mice were placed in a resting home cage with new bedding for 30 min. Mice were then placed in a regular Plexiglas home cage (L, 254 mm × W, 152 mm × H, 127 mm) that contained a modified 50 mL Falcon tube (89 mm × 32 mm) in the center. Two 5 min trials were conducted with a 30 min inter-trial interval. Time spent sniffing the Falcon tube was measured; sniffing was recognized if a whisker touched the tube.

##### Open field test

Time spent in the margin of an open inescapable field (i.e., thigmotaxis) and horizontal locomotor activity were measured to analyze anxiety-related behavior and motor activity under a higher level of stress, respectively. The mice were then observed for 30 min. The time spent in the margin area (i.e., thigmotaxis) and distance travelled (cm) in the open field were analyzed.

##### Olfactory responses to social and non-social cues

This test was conducted in a modified home cage (L, 28.5 cm × W, 17.5 cm × H, 12.5 cm) that had been divided into a 19.5-cm-long compartment and a 9-cm-long compartment using a partition wall with a 5 cm (H) × 5 cm (W) opening; the illuminance was ∼430 lux. The test was conducted as previously described (*53*), with slight modifications. First, the mice were habituated to the apparatus for 15 min. A 1 mL Eppendorf tube with seven holes (one in the middle and six surrounding it) in the cap contained a filter paper scented with a test odor. The tube was attached to the cage wall using Velcro. The filter paper (#3698-325; Whatman, Maidstone, UK) was soaked in 10 µL of each odorant. The following odors were sequentially tested: water, almond, banana, urine from a 2-month-old non-littermate C57BL/6J male (C57-1), urine from another 2-month-old non-littermate C57BL/6J male (C57-2), and urine from the first C57BL/6J mouse (C57-1). Urine was collected before testing and frozen at −20 °C until the test day. Each odor was tested in three 2 min trials with 10 s inter-trial and inter-session intervals. The time spent sniffing the tube was measured and analyzed.

#### ChIP-seq

We conducted ChIP-seq using two slightly different protocols.

##### Protocol 1

###### Preparation of fragmented DNA

We used a culture of neural progenitor cells derived from the hippocampus of P0 C57BL/6J pups. Neural stem/progenitor cells derived from hippocampal tissue from each pup were separately cultured. Cells from the four pups were pooled to make one sample (1 × 10^7^ cells per tube). The four pups were derived from the same mother, and three sets of four pups were used. Cells were collected by centrifugation (500 ×*g*, 5 min at 25 °C), resuspended in 10 mL PBS, washed (500 ×*g*, 5 min at 25 °C), and resuspended in culture medium (9 mL). Formaldehyde (270 µL, 37%, S25329; Fisher, Fair Lawn, NJ, USA) was added and mixed for cross-linking. Cells were incubated for 10 min at 25 °C on a rocking platform, and formaldehyde was quenched by glycine (1 mL, 1.25 M, BP381-500; Fisher). Cells were collected by centrifugation (500 ×*g*, 5 min at 25 °C) and washed with ice-cold PBS (10 mL) three times. Cells were re-suspended in 1 mL lysis buffer (CHP1, Imprint Chromatin Immunoprecipitation Kit; Sigma) and 10 µL proteinase inhibitor cocktail (1% SDS, 50 mM Tris-Cl, pH 8.1, 10 mM EDTA; Sigma) and incubated on ice for 1 h. DNA was sheared on ice by sonication with a Bioruptor four times (level: H, cycle: 30 s, 15 m) and centrifuged at 20,817 ×*g* for 10 min at 4 °C. The supernatant was transferred to a new tube and stored at –80 °C. The average fragment size was 200–400 bp.

###### Chromatin immunoprecipitation

We used the Imprint ChIP Kit (Sigma). We used sonicated cell lysates (1 × 10^6^ cell-derived sample per well). Cell lysates (100 µL) were diluted with 100 µL antibody buffer. Diluted cell lysates were stored as input DNA samples. Normal IgG (1 µg, M8695, Sigma) and anti-TBX1 antibody (1 µg, #109313; Abcam) were also diluted with 100 µL antibody buffer and added to strip wells. Strip wells were covered with Parafilm (PM-999; Pechiney Plastic Packaging, Chicago, IL, USA) and placed on a shaker for 60–90 min at 25 °C. The wells were washed thrice with antibody buffer (150 µL). Diluted cell lysates (200 µL DNA/well) were added to each well. Wells were covered with parafilm again and incubated at 25 °C on a shaker for 60–90 min. The supernatant was removed, and the wells were washed six times with 150 µL IP wash buffer, allowing 2 min for each wash. Wells were washed once with 150 µL 1× Tris-EDTA buffer. For cross-link reversal, DNA release buffer (100 µL) and protease K (2.5 µL) were added to the wells. The wells were covered with stripcaps and incubated at 65 °C for 15 min. Reversing solution (100 µL) was added to the samples, recovered, and incubated at 65 °C for 90 min. The samples were then transferred to new tubes.

###### DNA purification

Next, 250 µL TE-saturated phenol (#15513-039 GIBCO BRL; Thermo Fisher) and 250 µL chloroform (C298; Fisher) were added to both the chromatin immunoprecipitated samples and the input DNA samples and vortexed. After a 15 min incubation at 25 °C, the samples were centrifuged at 17750 x*g* (14,000 rpm) for 10 min at 25 °C. The supernatant was transferred to a 1.5 mL Eppendorf tube; next, 10 µL 3M sodium acetate and 250 µL 100% EtOH were added, vortexed, and incubated for 1 h at –20 °C. Samples were centrifuged at 20380 x*g* (15,000 rpm) for 20 min at 4 °C. The supernatant was discarded, 200 µL 70% EtOH was added, and the mixture was centrifuged at 17750 x*g* (14,000 rpm) for 10 min at 4 °C. The supernatant was discarded, and the pellet was dried and resuspended in 20 µL Tris-EDTA. Preliminary screening compared the enrichment of *Pten* promoter segments relative to normal IgG-precipitated samples and non-precipitated samples (Input). The data showed robust enrichment of signals in TBX1-precipitated samples compared to input and normal (5%) precipitated samples. The prepared samples were analyzed at the Epigenomic Core of the Albert Einstein College of Medicine.

###### ChIP-seq

The DNA was ligated using Illumina adapters (San Diego, CA, USA) with limited PCR amplification to create the Illumina library. Enrichment was confirmed by quantitative PCR (qPCR). A library was created using the input DNA sample as a control. The ChIP and input control libraries were sequenced simultaneously using Hiseq2500. This involved the first step of cluster generation *in situ* on the flow cell, followed by massively parallel sequencing using fluorescently labeled reversible terminator nucleotides. The clustered templates were sequenced based on the bases during each read. Images generated by Hiseq2500 were processed using the Hiseq Sequence Control Software (HCS v2 with RTA v1.17, and CASAVA v1.8). These software components perform the functions of image analysis (Firecrest), base-calling (Bustard), and alignment of sequence tags (up to 100 bp) to the appropriate reference genome (ELAND). Throughout this process, quality metrics were recorded to measure the experimental efficacy and to facilitate rigorous filtering of sequences prior to genome alignment. CisGenome software was used to identify protein-binding positions from the aligned sequence tags. This software first performed a sliding window exploratory analysis to estimate the false discovery rate (FDR) by comparing sequence counts for ChIP and input samples within each window, generating a conditional binomial distribution for model enrichment. Peak detection was then performed to predict the binding sites given a specified FDR threshold. Peaks were visualized as tracks alongside relevant genomic annotation tracks on the UCSC genome browser (mm10).

Given a specific stringency, a set of statistically relevant peaks was identified. The location and level of each peak in the genome, as well as the association of peaks with different genomic features, were described using summary statistics and standard modeling approaches. Comparison of peak sets across independent samples was conducted by aligning peaks and testing for over-representation in a particular class of samples using a window-based approach. The signal peaks were aligned with respect to the known genes. The reads were created using the Integrative Genomics Viewer (IGV) v2.3.59. The binding sites were classified in relation to transcription start sites using GREAT v4.0.4 (*54*).

##### Protocol 2

###### ChIP

We used a culture of neural progenitor cells derived from the hippocampus of P0 C57BL/6J pups. Cells from four pups were pooled to make one sample (4 × 10^6^ cells per IP prep). The four pups were derived from the same mother, and four sets of four pups were used. ChIP was performed using the SimpleChIP® Enzymatic Chromatin IP Kit (#9005S; Cell Signaling Technology, Danvers, MA, USA) according to the manufacturer’s protocol. All buffers, reagents, and DNA spin columns were included in the kit unless otherwise stated. Briefly, cells were cross-linked with 1% formaldehyde (37%, F8775; Sigma-Aldrich) in 10 mL culture medium for 10 min and quenched with 1 mL 10× glycine. Cells were collected by centrifugation (500 ×*g*, 5 min at 4 °C) and washed twice with 20 mL ice-cold PBS. Cells were re-suspended in 1 mL ice-cold Buffer A containing DTT and protease inhibitor cocktail and incubated on ice for 10 min. Nuclei were pelleted by centrifugation (2,000 ×*g*, 5 min at 4 °C) and re-suspended in 1 mL ice-cold Buffer B containing DTT. Micrococcal nuclease (0.5 µL per IP prep) was added to digest chromatin and incubated for 20 min at 37 °C. Nuclei were collected by centrifugation (1,600 ×*g*, 1 min at 4 °C) and re-suspended in ChIP Buffer containing protease inhibitor cocktail, and then incubated for 10 min on ice. Nuclear membranes were broken by sonication (20 s, on ice, thrice). The samples were centrifuged (9,400 ×*g*, 10 min at 4 °C), and the supernatant was stored at –80 °C. Each supernatant (50 µL chromatin sample) was purified (described below), and the DNA concentration was determined. DNA fragment sizes were determined by electrophoresis, and the sizes were 150–900 bp.

Cell lysates were diluted with ChIP Buffer containing protease inhibitor cocktail, and 10 µg chromatin DNA in 500 µL ChIP buffer was used per IP prep. Cell lysates were stored as input DNA samples. Negative control normal rabbit IgG (2 µg) or anti-TBX1 antibody (2 µg, ab109313, lot# YH080909D; Abcam) was added to the lysate and incubated overnight at 4 °C with rotation. ChIP-Grade Protein G Magnetic Beads (30 µL per sample) were added to each tube and incubated for 2 h at 4 °C with rotation. The beads were washed with 1 mL low-salt wash buffer in a magnetic separation rack thrice and then with 1 mL high-salt wash buffer. ChIP elution buffer (150 µL per sample) was added to each tube, and chromatin samples were eluted from the beads by incubation for 30 min at 65 °C with rotation. Protease K (2 µL) and 5 M NaCl (6 µL) were added to each sample, and the samples were incubated for 2 h at 65 °C to reverse cross-linking.

###### DNA purification

DNA-binding buffer (750 µL) was added to each ChIP-DNA sample and transferred to a DNA spin column. The columns were washed with 750 µL DNA washing buffer, and the purified ChIP-DNA samples were eluted with 50 µL DNA elution buffer from the columns.

###### ChIP-seq

The purified DNA samples were submitted to Arraystar (http://www.arraystar.com/) for library construction, sequencing, and basic data analyses. Illumina genomic adapters were used to prepare the ChIP-seq library. Libraries were quantified using an Agilent 2100 Bioanalyzer. The libraries were denatured with 0.1 M NaOH to generate single-stranded DNA molecules, captured on an Illumina flow cell, and amplified *in situ*. The libraries were then sequenced on an Illumina NovaSeq 6000 using the NovaSeq 6000 S4 reagent kit (300 cycles).

After sequencing was performed using the sequencing platform, image analysis and base calling were performed using the Off-Line Basecaller software (OLB v1.8; Illumina). After the Solexa CHASTITY quality filter was passed, the clean reads were aligned to the mouse genome (UCSC MM10 using BOWTIE v2.1.0). Aligned reads were used for peak calling of ChIP regions using MACS v1.4.0. Statistically significant ChIP-enriched regions (peaks) were identified by IP using a *p*-value threshold of 10^-4^. The peaks in the samples were annotated by the nearest gene using the newest UCSC RefSeq database.

The UCSC Genome Browser GRCm38/mm10 (http://genome.ucsc.edu/) was used to plot the genomic positions of the peak regions (*55*). MoLoTool (http://molotool.autosome.ru/) was used to predict the T-binding motif (*56*) (see **Fig. S12**, **S16**, **S17**, and **S18**).

###### Gene annotation

The genomic locations described in the files from the DBD databases were used to generate BED files for detailed annotation based on GRCh38 (hg38). The BED files were converted to GRCh38 (hg38) using UCSC LiftOver (*57*) with a 95% minimum ratio of bases that must be remapped. Annotation was performed using a combination of ChIPSeeker (*58*) and biomaRt R package (*59*). The files from MANGO, SFARI, and DDG2P did not contain genomic locations; therefore, the gene symbols were used for annotation with biomaRt.

The networks were constructed using the STRING database (*41*); gene lists from MANGO, DBD, and SFARI; and a pooled gene list with DBD and SFARI. The graphs were plotted using the DiagrammeR^2^ R package (*60*) and the resulting edge lists from STRING with a combined score threshold of 0.4 and edge width corresponding to the combined score.

#### ChIP-PCR

We investigated whether endogenous TBX1 protein occupies the TBX1 transcriptional response element (TRE) of *Pten* in postnatal neural progenitor cells derived from the P0 hippocampal dentate gyrus of C57BL/6J mice. A pair of primers was designed to detect a sequence that contains the TBX1-binding sites up to 1 kb upstream of *Pten* (**Fig. 5B**). We carried out the UCSC In-Silico PCR and found that this pair did not detect any promoter other than that of *Pten*. The purified DNA samples were prepared using the same sample preparation protocol used for ChIP-seq. Cells (4 × 10^6^) were used for immunoprecipitation. qPCR was performed with the positive control histone H3 sample, the negative control normal IgG sample, and a serial dilution of the 2% input chromatin DNA (undiluted, 1:5, 1:25, 1:125) to create a standard curve. The percent input was calculated as follows: Percent Input = 2% × 2(C[T] 2%Input Sample − C[T] IP Sample). C[T] = CT = threshold cycle of PCR. The assay was performed five times.

#### Luciferase assay

We generated several promoter sequence constructs of murine PTEN (see **Table S1-1**; **Figure S14**) using our published procedure (*23*) with a slight modification (#638909, In-Fusion HD Cloning Plus; Clontech, Mountain View, CA, USA). Promoter cloning was conducted using murine *Pten* mRNA (-201) (Ensemble genome browser). An approximately 1 kb segment of promoter containing putative T-box–binding sequences was cloned into the multiple cloning site of a cloning vector, using standard restriction-enzyme cloning or Gibson Assembly Master Mix (E2611; New England Biolabs, Ipswich, MA, USA). All constructs were verified by Sanger sequencing of plasmids. Fragments, taken using the 1 kb promoter sequence, were infused into pMettLuc-2 Reporter (PT4059-5; Clontech) by *Eco*R1 (R0101; New England Biolabs) and amplified. The following additional sequences were also included for infusion: forward primer, 5’-tcaagcttcgaattc-3’; reverse primer, 5′-gtcgactgcagaattc-3’. Bands were excised using a Qiaquick Gel Extraction Kit (#28704; Qiagen).

We also used the QuickChange Site-Directed Mutagensis Kit (#200518; Agilent Technologies, Santa Clara, CA, USA) to induce mutations within the segment identified to be critical for the transcriptional regulation of the PTEN promoter, following the manufacturer’s instructions.

The transcriptional activity of endogenous TBX1 was examined using a luciferase reporter assay (Ready-To-Glow™ secreted luciferase reporter system, #631726; Clontech). P0 neural progenitor cells of the hippocampus (2 × 10^5^ cells/well in a 6-well plate, 37 °C overnight) were transfected with pMetLuc2-promoter reporter vector (DMEM/F12+B27+N2+20 ng/mL EGF) in Opti-MEM (#31985062; Invitrogen) and Lipofectamine LTX Plus (#15338100; Invitrogen) under our standard cell culture protocol (*11*) (see also “Cell culture” above). Luciferase activity was significantly higher with EGF than without it (**Figure S13**). The pMetLuc2 reporter plasmid was used as a negative control, and the pMetLuc2 control plasmid was used as a positive control. PCR with KOD-Plus-Neo (KOD-401; Toyobo, Osaka, Japan) was performed with 35 cycles at 94 °C for 2 min, 98 ℃ for 10 s, and 68 ℃ for 30 s. The pSEAP2-control vector was used as an internal control to normalize the cell numbers between groups. The culture medium was collected 48 h post-transfection, and the luciferase activity secreted in the culture medium was measured in triplicate in 96-well plates using the Synergy4 Microplate Reader (S4MLFPTA; BioTek, Winooski, VT, USA). The plasmid with the full promoter sequence had ∼30-fold higher activity than that with no promoter (**Figure S14**; *P* = 0.0022). Relative luciferase activity was calculated by normalizing the results to the samples with the full-length promoter sequence.

#### qRT-PCR for validation of TBX1’s target genes

Total RNA was extracted using an RNeasy Plus Mini Kit (#74134; Qiagen, Germantown, MD, USA), according to the manufacturer’s instructions. cDNA was synthesized from total RNA using SuperScript IV VILO Master Mix (#11766050; Invitrogen). qPCRs were performed in triplicate on a QuantStudio 6 Flex Real-Time PCR System (#4485694; Applied Biosystems, Waltham, MA, USA) using the Power SYBR™ Green PCR Master Mix (#4367659; Applied Biosystems). Data were analyzed using the ΔΔCt method and normalized to the reference gene Cyc1 (SY-mo-600; PrimerDesign, Chandler’s Ford, UK).

#### Statistical analysis

Data from two or more groups were compared using Student’s *t*-test or analysis of variance (ANOVA), respectively. Newman–Keuls post-hoc comparison was used when an interaction was significant. Normality and variance homogeneity were evaluated using the Shapiro–Wilk test and Levene’s homogeneity of variance test, respectively. When either assumption was violated, data were analyzed using a generalized linear mixed model or via Mann–Whitney or Wilcoxon tests. The minimum significance level was set at 5%. As multiple comparisons were limited to the genotype or treatment factor at different levels of the other factor and not applied to every possible comparison (i.e., planned comparison), we did not apply correction.

---

**Figure S1.**
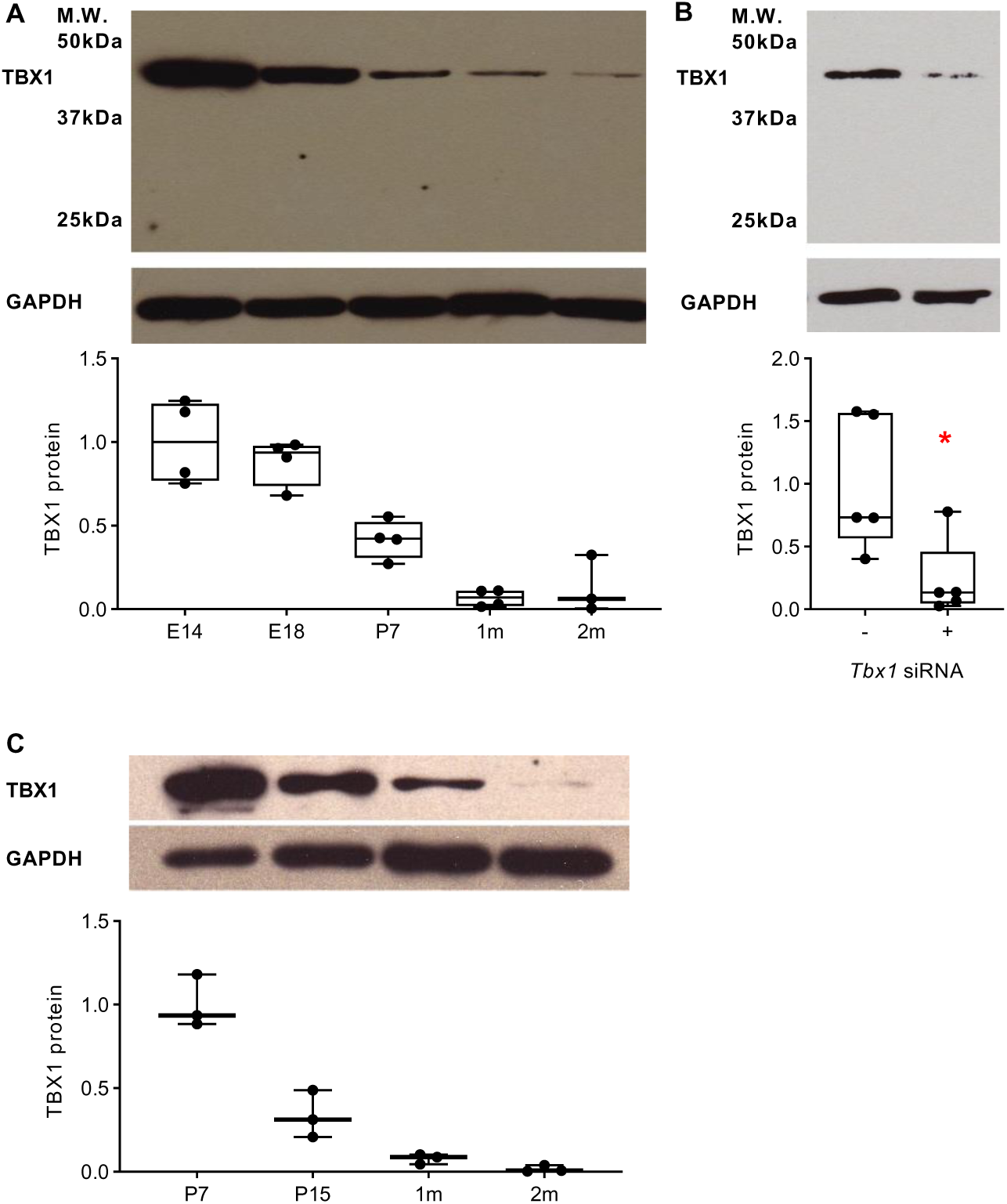
Developmental changes in TBX1 in the C57BL/6J mouse brain. (**A**) TBX1 protein levels in the whole brain declined from the embryonic to adult periods (*P* < 0.001). Whole-brain samples of C57BL/6J mice were prepared from embryonic day 14 (E14; *N* = 4), E18 (*N* = 4), postnatal day 7 (P7; *N* = 4), 5-week-old (1m; *N* = 4), and 8-week-old (2m, *N* = 3) mice. (**B**) The specific TBX1 band was identified using cell culture of P0 stem/progenitor cells derived from the P0 hippocampus of C57BL/6J mice (\**P* = 0.043). Neural stem/progenitor cells received either control siRNA (siGLO RISC-Free Control siRNA) or *Tbx1* siRNA (40 nM, Santa Cruz Biotechnology, Santa Cruz, CA, USA) for 72 h. Total *N* = 5 per group. (**C**) TBX1 levels declined during the post-embryonic period in the hippocampus of C57BL/6J mice (*P* < 0.001). Hippocampal samples (10 µg) were prepared from P7, P15, 1m, and 2m mice. *N* = 3 per age. TBX1 protein was calculated by dividing the intensity of TBX1 signals by that of GAPDH; the average intensity of E14 (**A**), no *Tbx1* siRNA (**B**), or P7 (**C**) was set as 1.0. All values are shown in box-and-whisker plots. M.W., molecular weight; GAPDH, glyceraldehyde-3-phosphate dehydrogenase.

**Figure S2.**
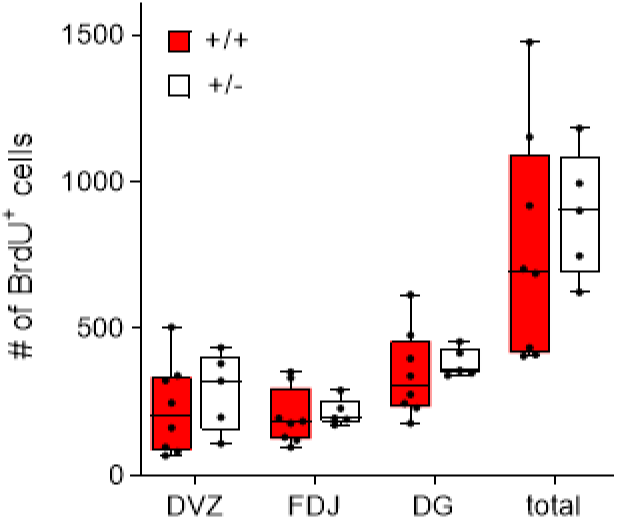
Embryonic neurogenesis in the hippocampus of *Tbx1*^+/+^ and *Tbx1*^+/-^ mice. The number (mean + SEM) of BrdU-positive cells in the embryonic hippocampus of *Tbx1*^+/+^ and *Tbx1*^+/-^ mice. BrdU was injected at E18.5, and embryos were sacrificed 2 h later. BrdU-positive cells in the hippocampal dentate gyrus (DG), dentate ventricular zone (DVZ), and fimbrio-dentate junction (FDJ) were indistinguishable between *Tbx1*^+/+^ and *Tbx1*^+/-^ mice (genotype, *P* = 0.5541; genotype × position, *P* = 0.574); the total number of BrdU-positive cells in all the three regions was indistinguishable between *Tbx1*^+/+^ and *Tbx1*^+/-^ mice (*P* = 0.554). Three sections per animal were counted. +/+, *N* = 8; +/-, *N* = 5.

**Figure S3.**
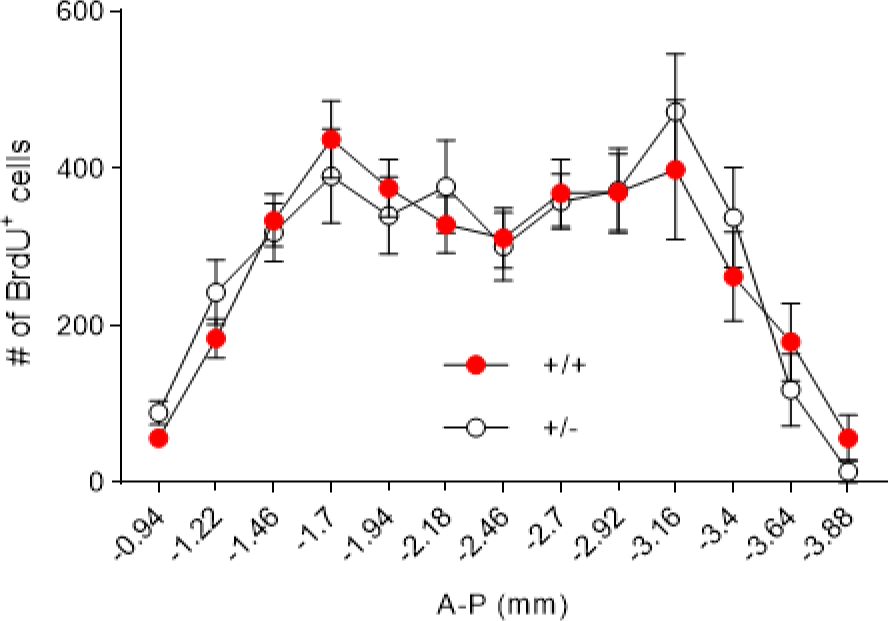
Adult neurogenesis in *Tbx1^+/+^ and Tbx1^+/-^* mice. The number (mean +SEM) of BrdU-positive cells in the hippocampus of *Tbx1^+/+^ and Tbx1^+/-^* mice at 5 weeks of age. Mice were given a single injection of BrdU (50 mg/kg, i.p.) 24 h before sacrifice at 5 weeks of age. (**A**) Number of BrdU-positive cells along the anterior-posterior axis in the hippocampal granule cell layer of *Tbx1*^+/+^ and *Tbx1*^+/-^ mice at 5 weeks of age at each anterior-posterior position (from the bregma, AP, −0.94 to −3.88) (genotype, F(1,15) = 0.016, *P* = 0.900; genotype × A-P position, F(12,180) = 0.684, *P* = 0.765). *Tbx1^+/+^*mice, *N* = 10; *Tbx1^+/-^* mice, *N* = 7.

**Figure S4.**
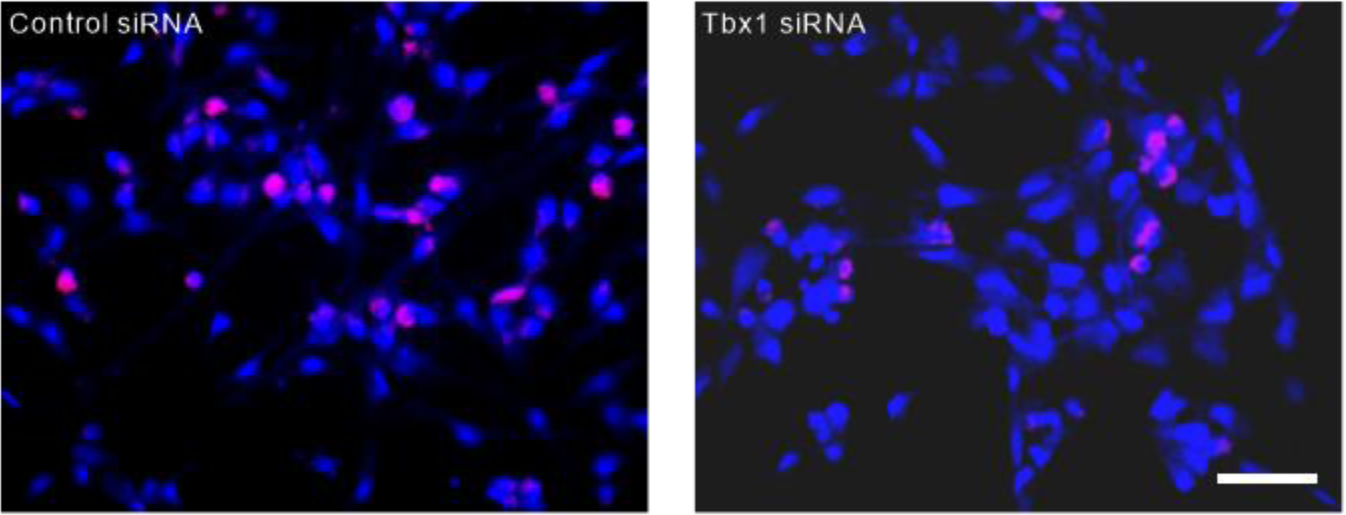
TBX1 is required for the proliferation of neonatal neural progenitor cells *in vitro*. Representative images of double-labeling of BrdU (red)- and DAPI (blue)-positive cells. *Tbx1* siRNA reduced the number of BrdU-positive cells (see **Fig. 2** for statistical analysis). Scale bar = 20 µm.

**Figure S5.**
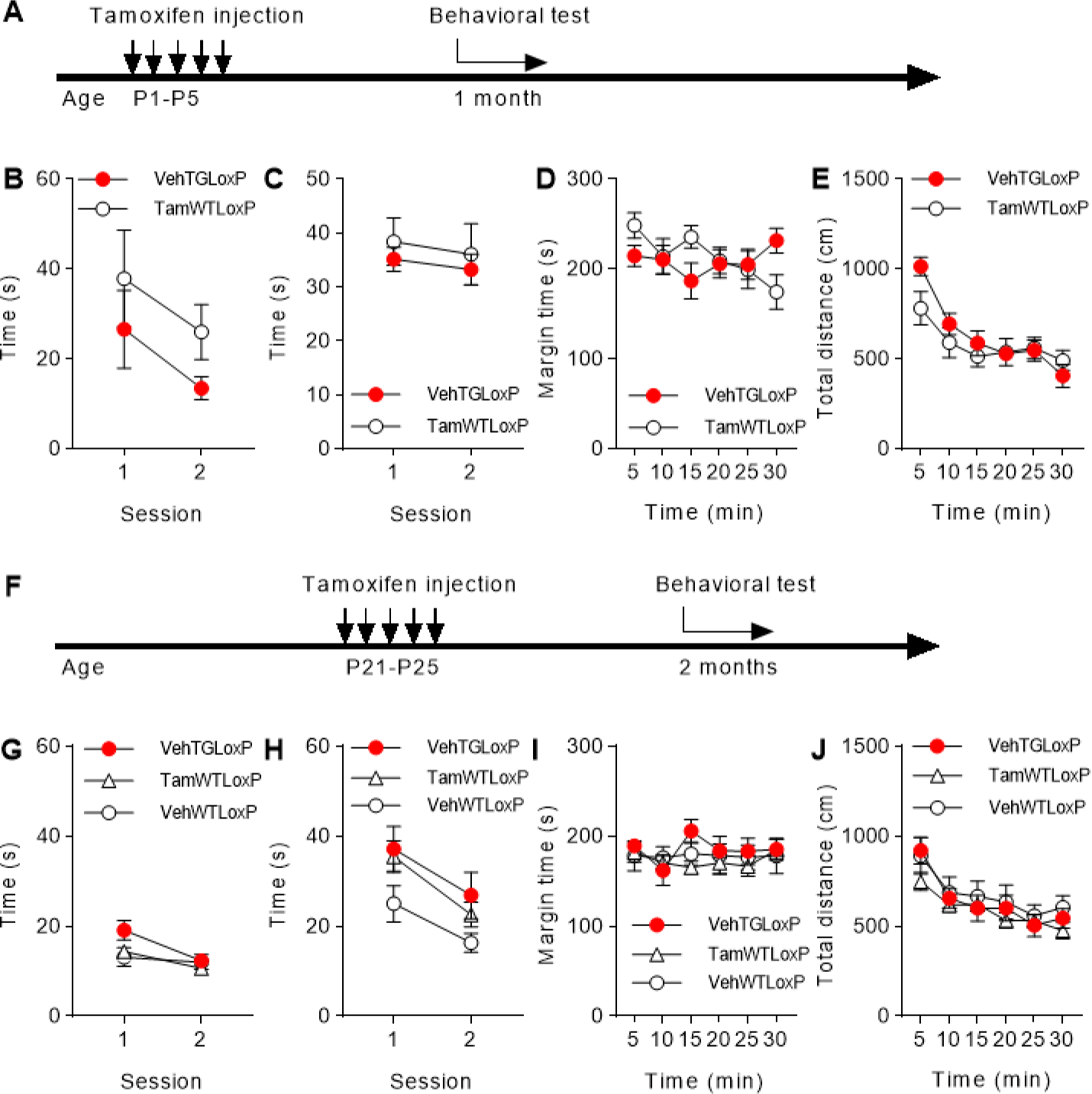
Behavioral phenotypes of mice with control genotypes. (**A**) Vehicle (Veh) or tamoxifen (Tam, 83.5 mg/kg body weight, i.p. via lactating mothers) was given to pups from P1 to P5, and behavioral assays started 1 month later. TGloxP included Nes-cre/ERT2;*Tbx1^flox/+^* and Nes-cre/ERT2;*Tbx1^flox/flox^* pups. WTLoxP included WT;*Tbx1^flox/+^* and WT;*Tbx1^flox/flox^* pups. These control groups did not differ in (**B**) social approach (TamWTLoxP, *N* = 12; VehTGLoxP, *N* = 7), (**C**) novel object approach (TamWTLoxP, *N* = 13; VehTGLoxP, *N* = 10), or (**D**) thigmotaxis and (**E**) locomotor activity (TamWTLoxP, *N* = 7; VehTGLoxP, *N* = 7) (genotype, *P >* 0.05 and genotype × time, *P >* 0.05 for all, except for **D** and **E:** genotype × time, *P* < 0.01). (**F**) Mice were directly injected with Veh or Tam (180 mg/kg, i.p. per day) from P21 to P25 and tested one month later (i.e., 2 months of age). These controls did not differ in (**G**) social approach (VehTGLoxP, *N* = 11; TamWTLoxP, *N* = 16; VehWTLoxP, *N* = 8), (**H**) novel object approach (VehTGLoxP, *N* = 10; TamWTLoxP, *N* = 16; VehWTLoxP, *N* = 7), or (**I**) thigmotaxis and (**J**) locomotor activity (VehTGLoxP, *N* = 12; TamWTLoxP, *N* = 19; VehWTLoxP, *N* = 9) (genotype, *P* > 0.05 and genotype × time, *P >* 0.05 for all).

**Figure S6.**
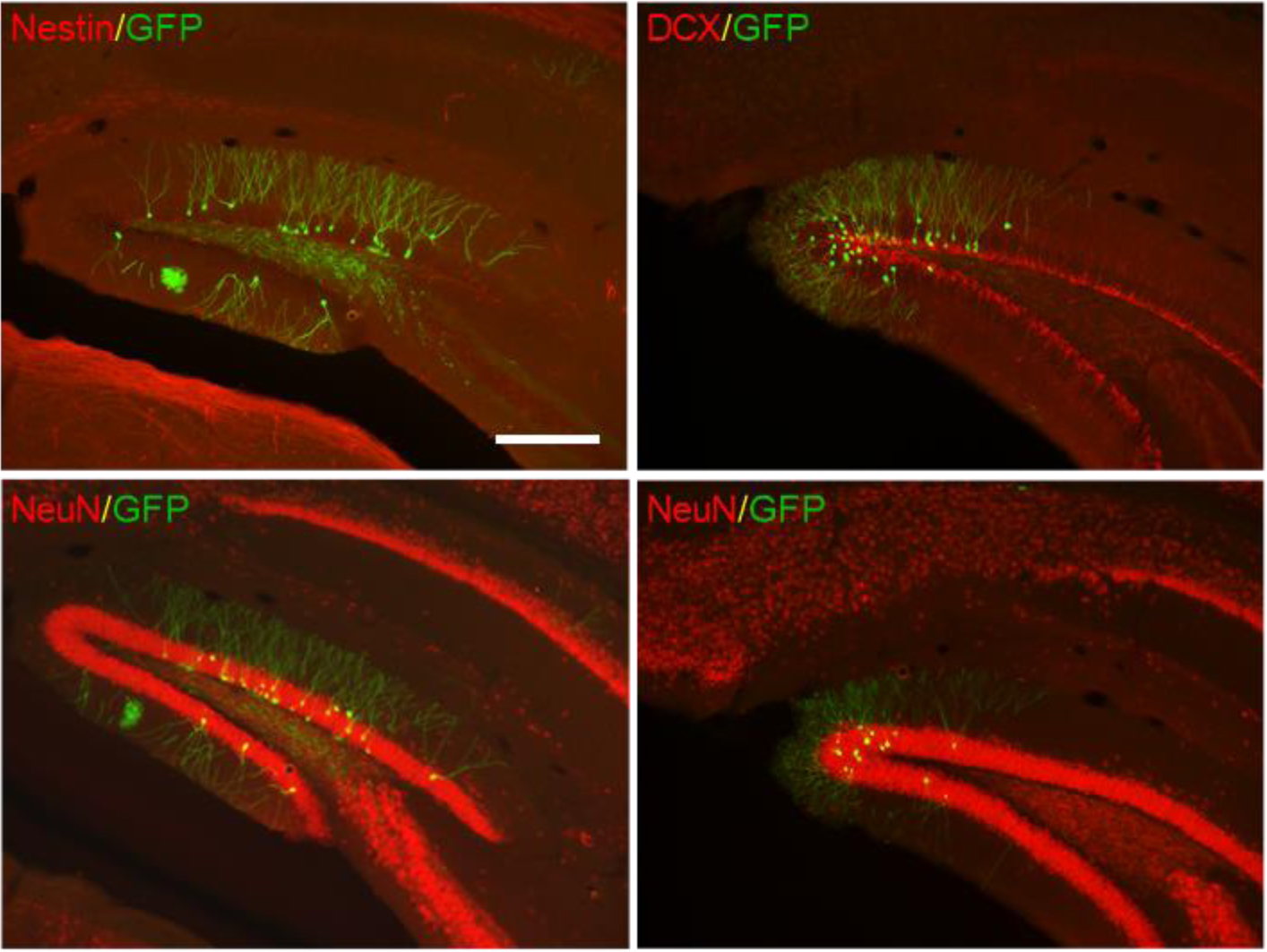
Recombination in tamoxifen-treated nestinCreERT;Ai3(RCL-EYFP) mice. Tamoxifen (83.5 mg/kg body weight to lactating mothers) was administered to pups from P1 to P5, and mice were sacrificed at 1 month of age. The selective expression of GFP-positive cells within the granule cell layer alone, lack of consistent GFP signals in brain regions with no adult neurogenesis, partial colocalization with DCX, and colocalization of GFP-positive cells in NeuN-positive cells only within the granule cell layer 1 month after tamoxifen were all consistent with the recombination that occurred selectively in stem/progenitor cells at P1–5. EYFP was stained with a GFP antibody. *N* = 3. Scale bar = 250 µm.

**Figure S7.**
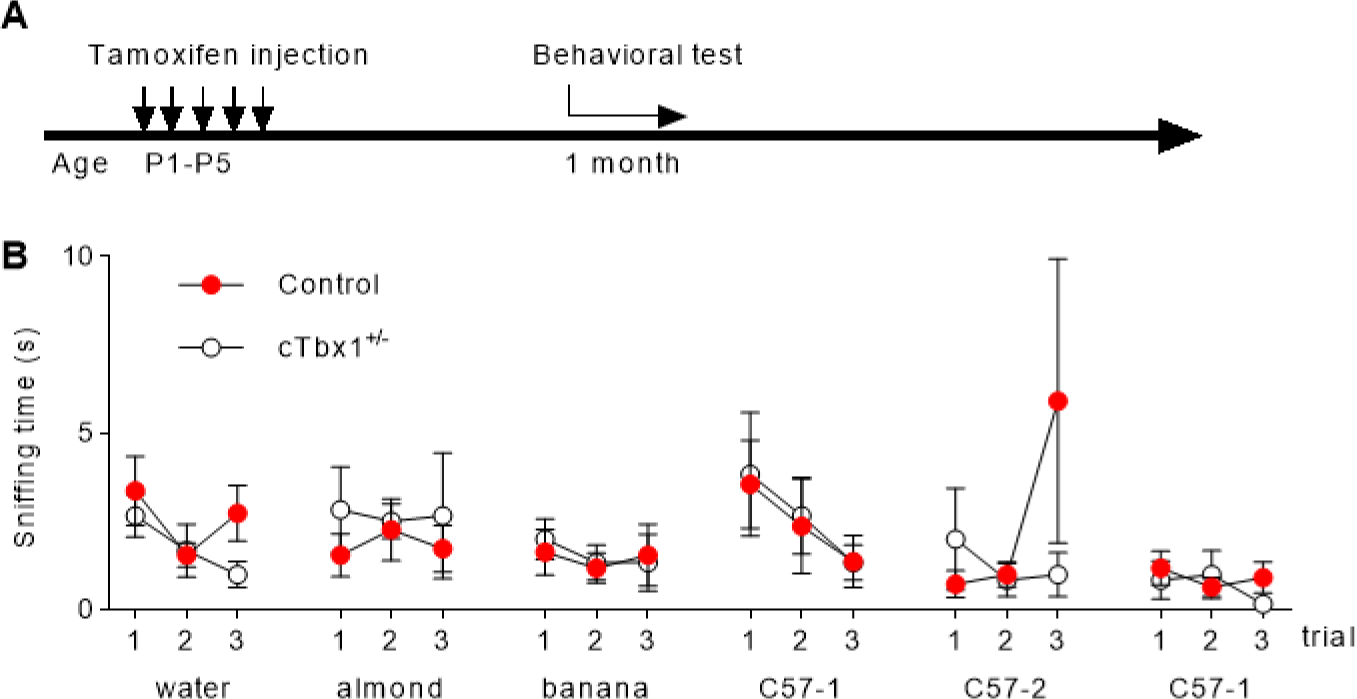
Effect of *Tbx1* deletion initiated during the neonatal (P1–5) period on olfactory responses of *cTbx1*^+/-^ mice. (**A**) Experimental design. (**B**) Olfactory response to various non-social and social odorants at 1 month of age following tamoxifen given at P1–5. Controls and *cTbx1*^+/-^ mice did not differ regardless of odorants or trials (genotype, F(1,15) = 0.065, *P* = 0.803; genotype × odorant, F(5,255) = 0.614, *P* = 0.689; genotype × trials, F(2,255) = 1.439, *P* = 0.239; genotype × trials × odorants, F(10,255) = 0.612, *P* = 0.803). The means ± SEM are shown. Control, *N* = 11; c*Tbx1*^+/-^, *N* = 6.

**Figure S8.**
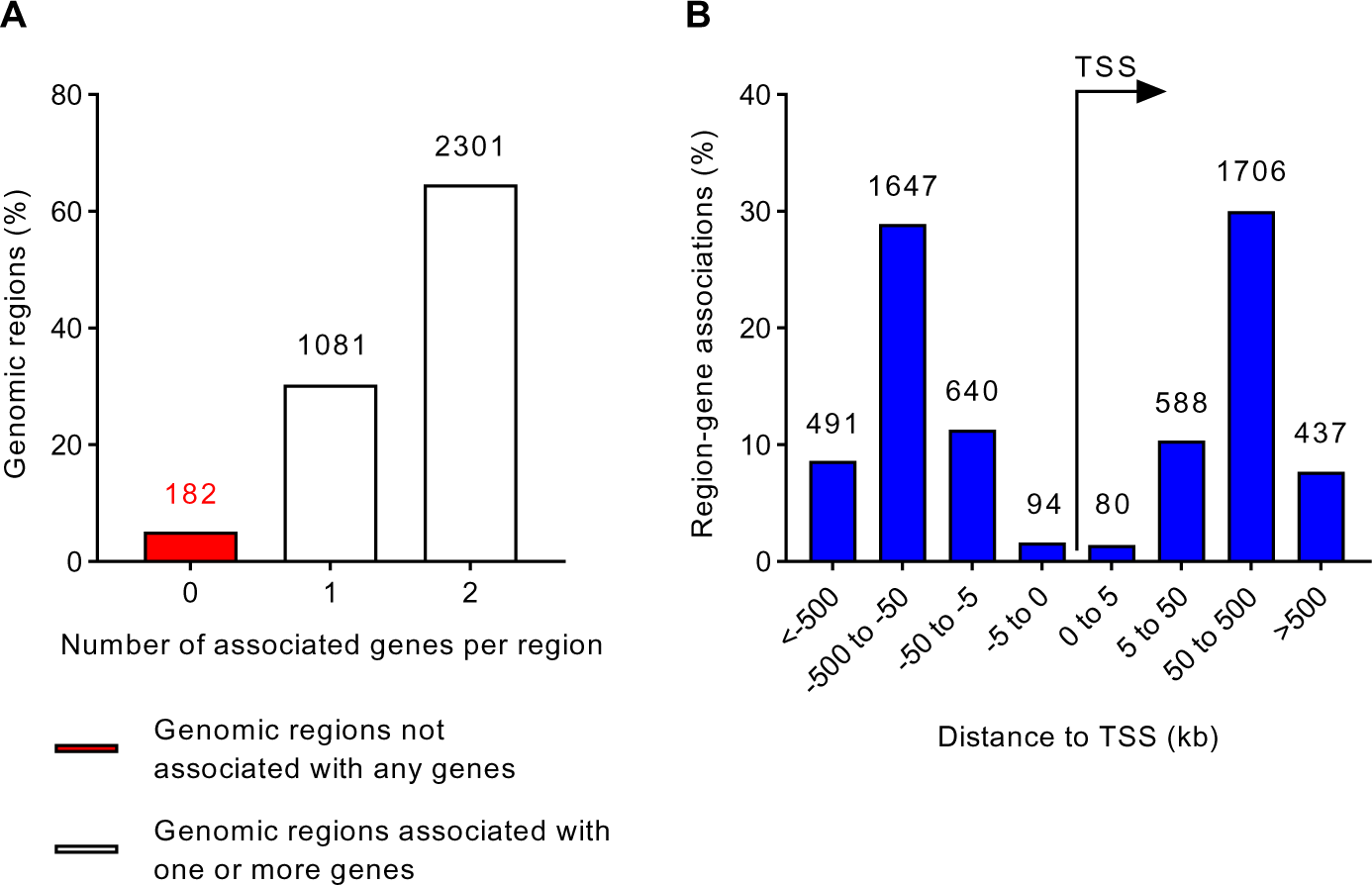
TBX1-binding sites in relation to the transcription start sites (TSSs). Data were analyzed based on GREAT v4.0.4 (http://great.stanford.edu/public/html/index.php).

**Figure S9.**
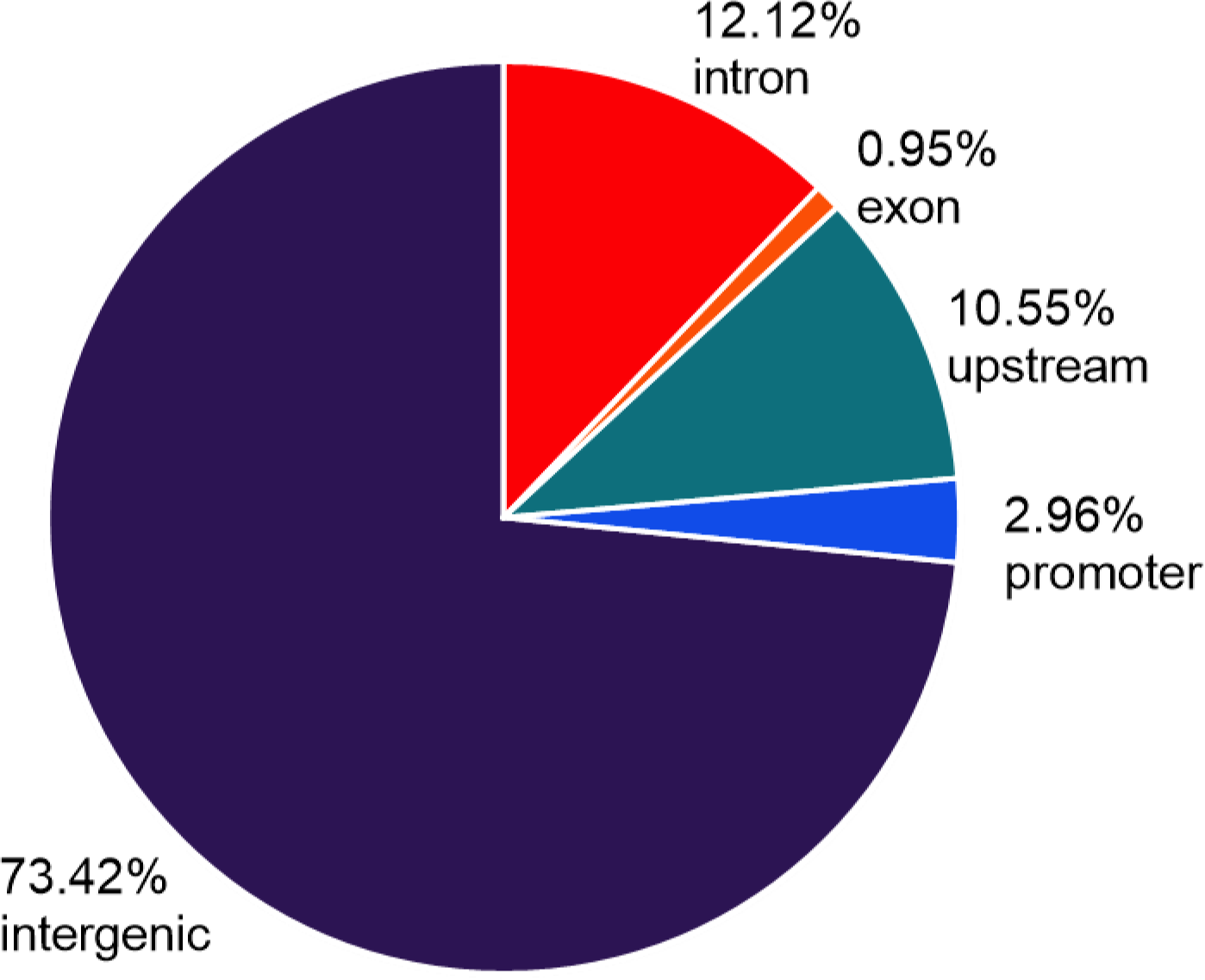
Proportions of TBX1’s binding peaks on the chromosome. Promoter, binding peaks are located 2,000 bp upstream and downstream from the transcription start site (TSS); upstream, binding peaks are located >2,000 bp to 20,000 bp upstream from the TSS; intron and exon, binding peaks are located in introns and exons, defined as starting at 2,000 bp downstream from the TSS; intergenic, peaks are located in none of the above.

**Figure S10-1.**
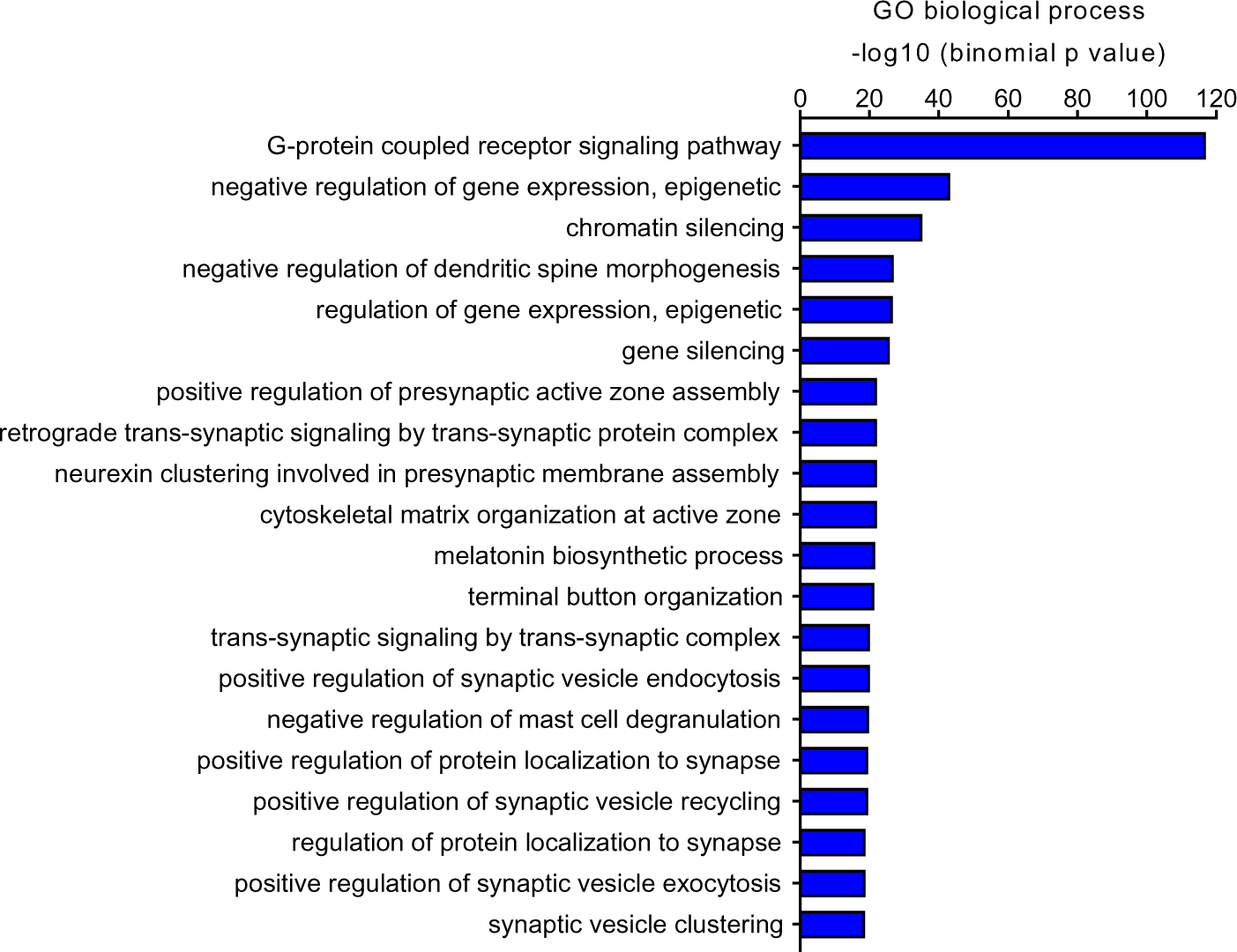
Biological processes associated with the target genes of TBX1.

**Figure S10-2.**
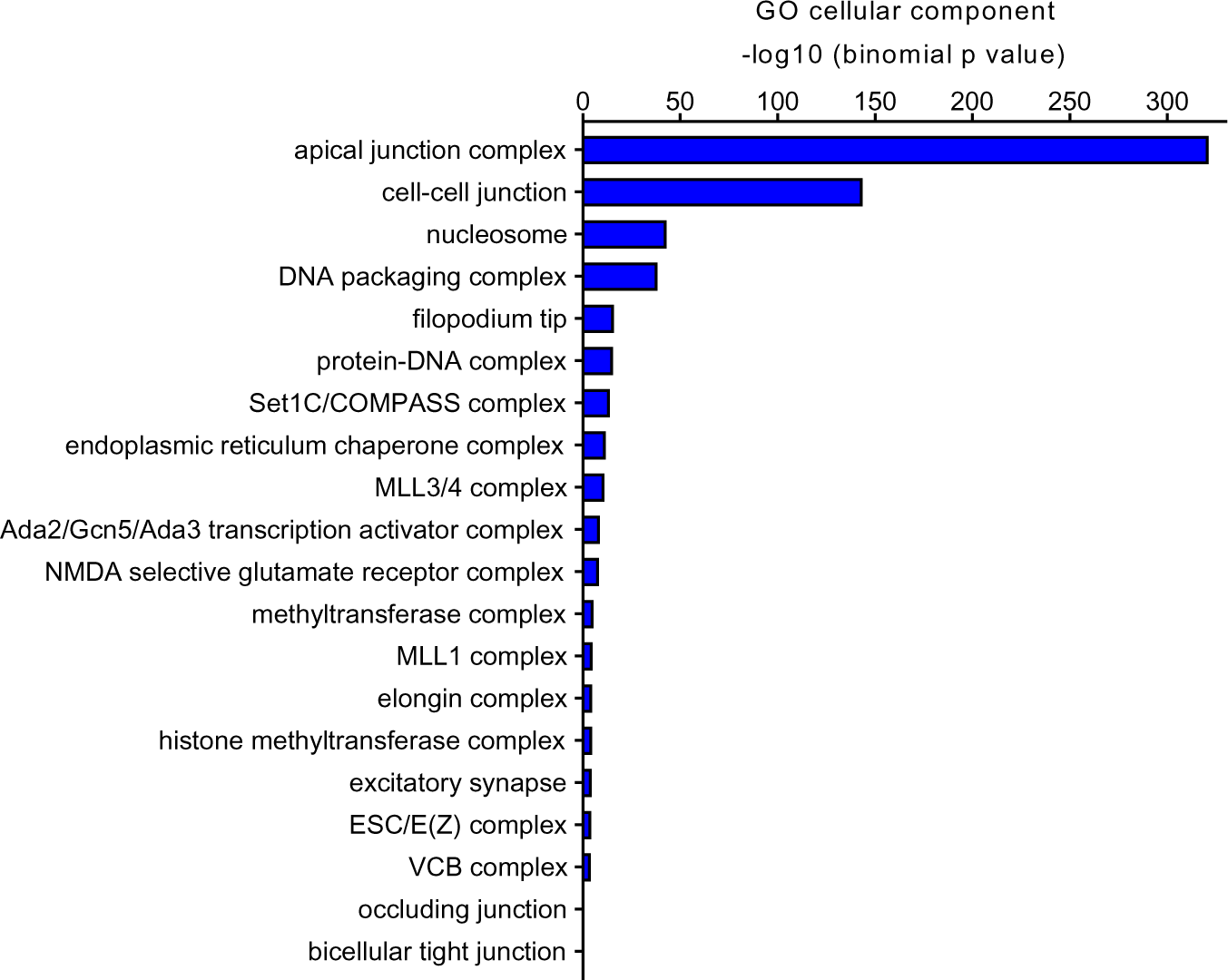
Cellular components associated with TBX1’s target genes.

**Figure S10-3.**
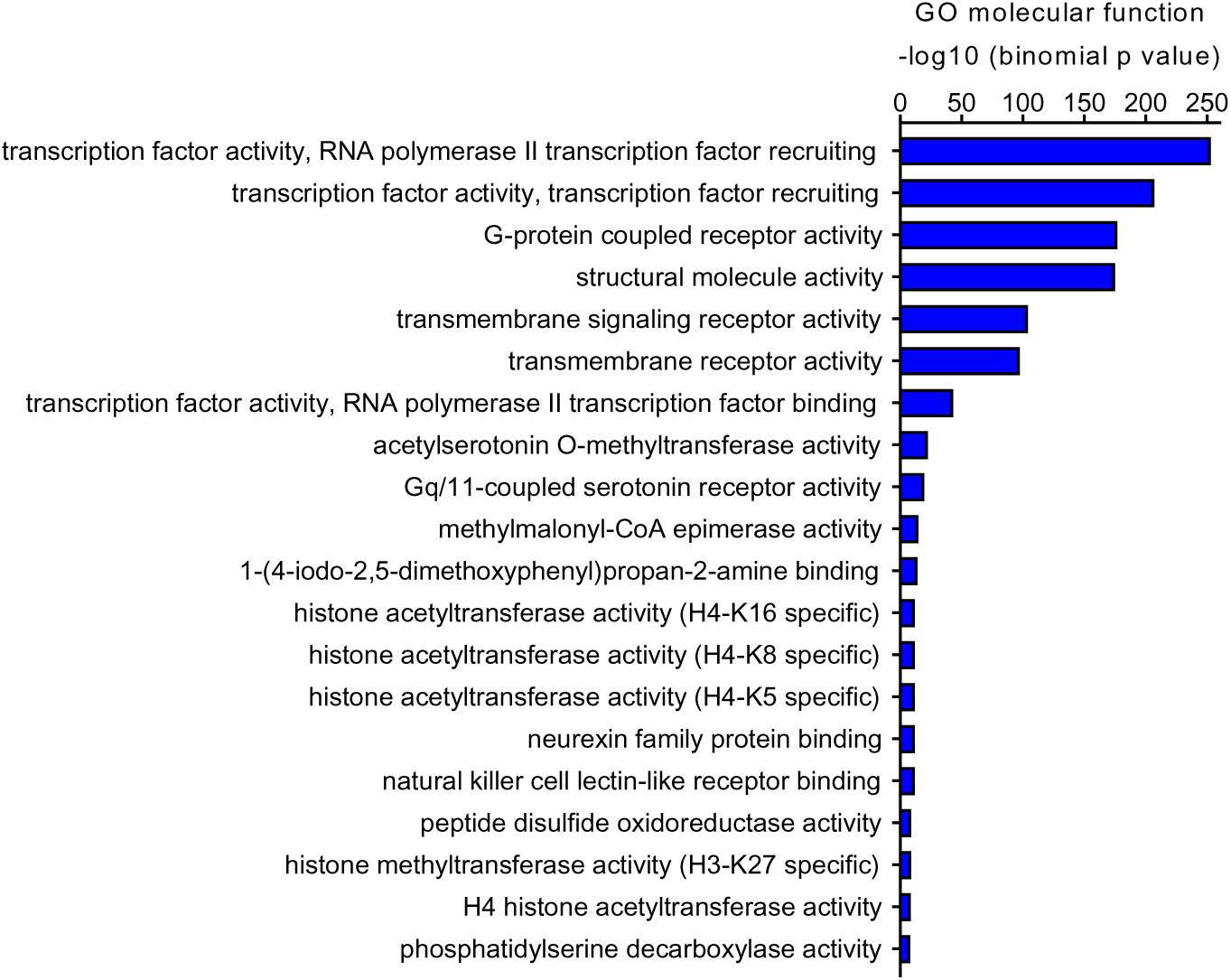
Molecular functions associated with TBX1’s target genes.

**Figure S11.**
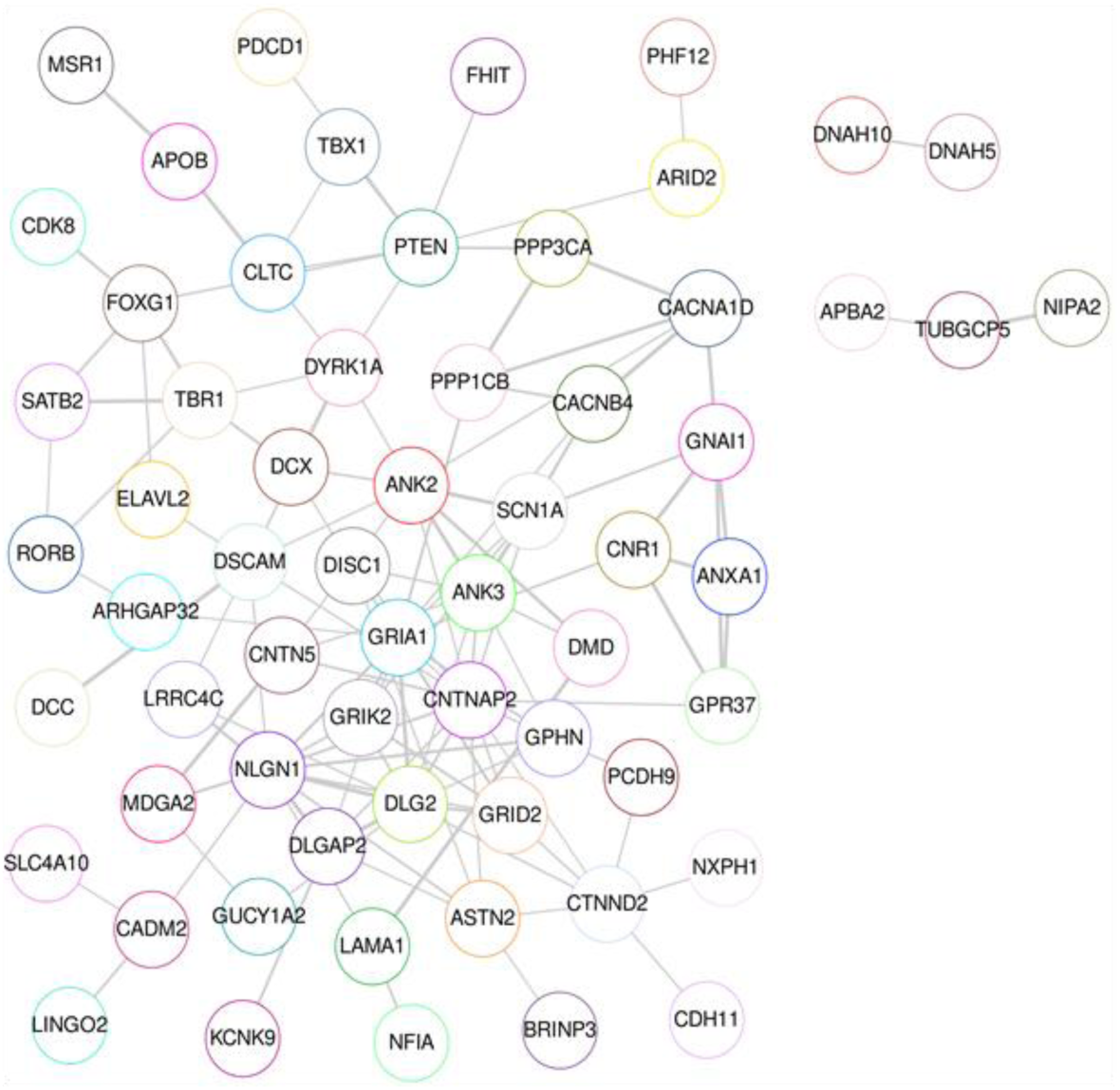
Potential interaction among genes bound by TBX1 searched in the gene databases of autism and neurodevelopmental disorders. SFARI (https://gene.sfari.org) and Geisinger Developmental Brain Disorder Database (https://dbd.geisingeradmi.org/) were used. Their potential interactions are shown, based on Search Tool for Retrieval of Interacting Genes/Proteins (STRING) analysis (*41*).

**Figure S12.**
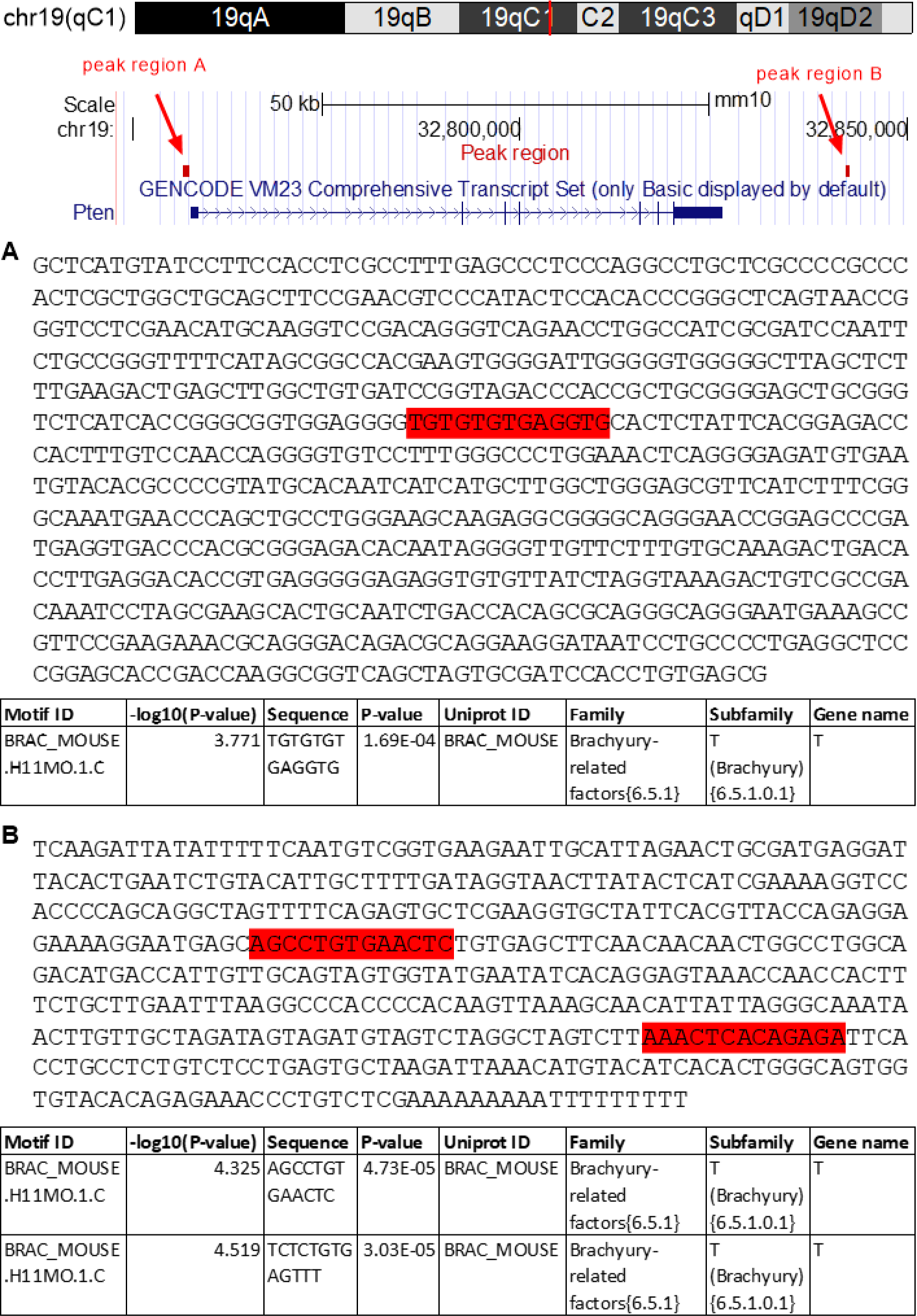
Two TBX1-binding sites near *Pten* and their nucleotide sequences (A and B).

**Figure S13.**
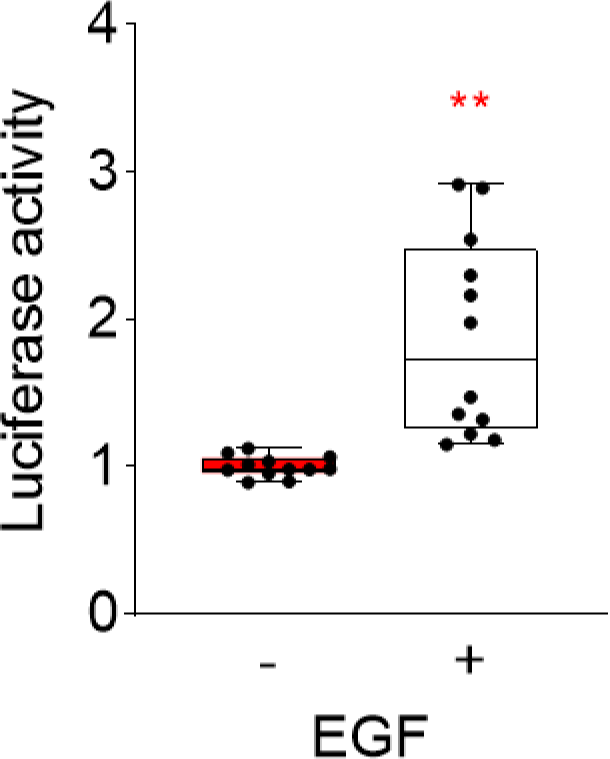
Promoter activity of *Pten*. Transcriptional activity of the 1 kb *Pten* promoter, as measured by luciferase activity, was higher with EGF (20 ng/mL) than without EGF (U = 144, *P* < 0.001; *N* = 12 per group). Twelve separate hippocampal tissue samples from two hemispheres were derived from six P0 C57BL/6J pups.

**Figure S14.**
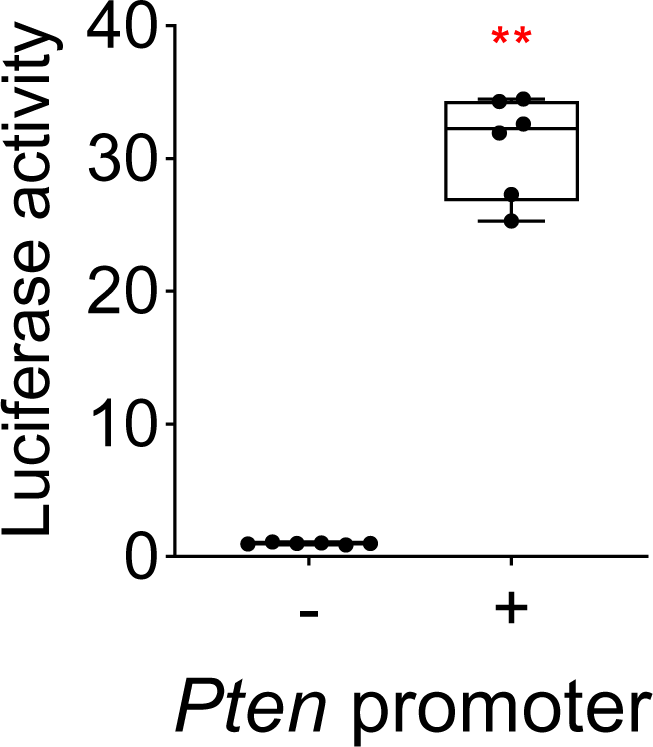
Promoter activity of *Pten*. Transcriptional activity of the pMetLuc2 reporter plasmid was significantly higher with than without the 1 kb full-length *Pten* promoter, as measured by luciferase activity (U = 144, *P* = 0.002; *N* = 6 per group).

**Figure S15.**
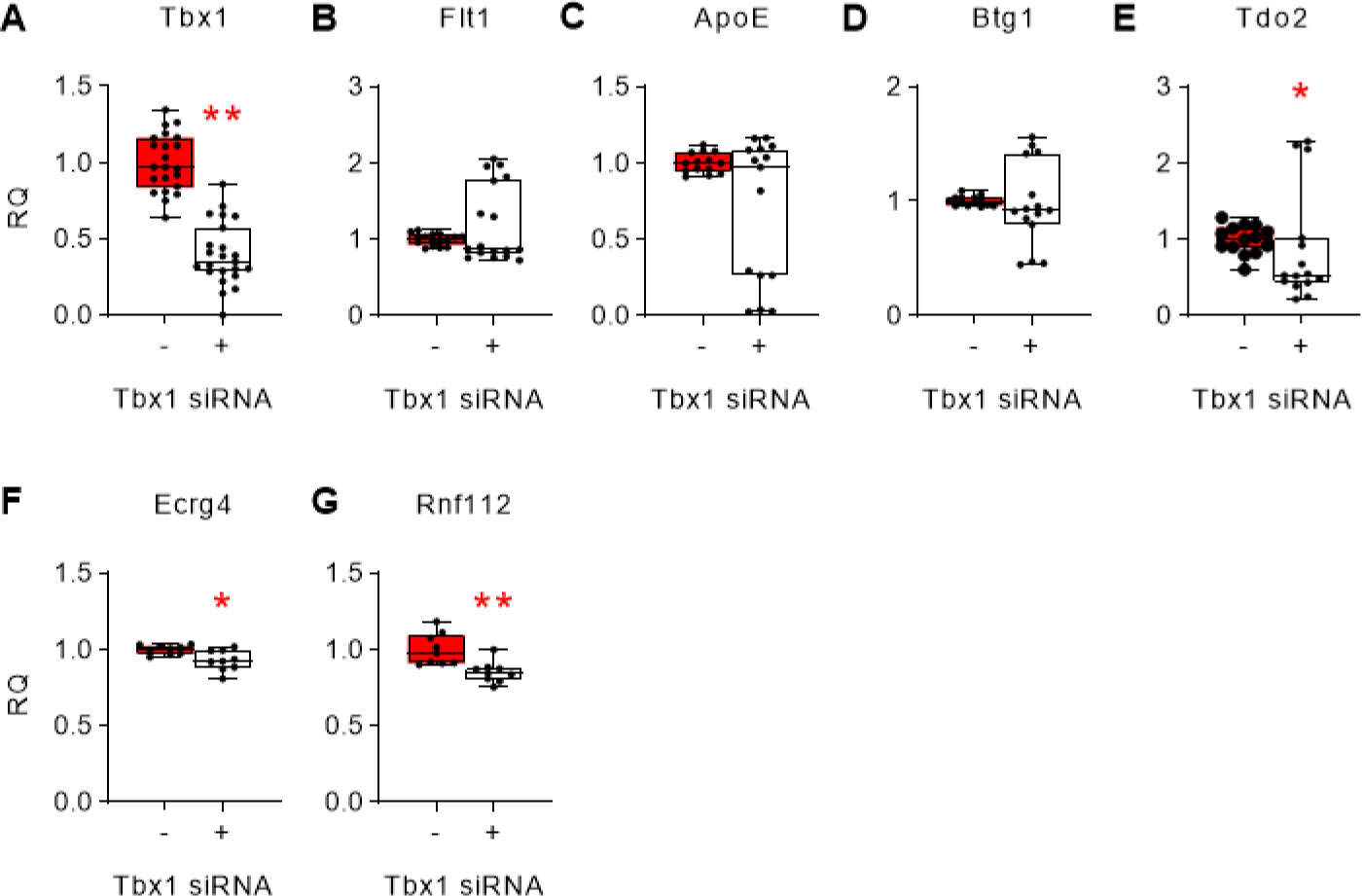
Effects of *Tbx1* siRNA on the mRNA expression levels (RQ) of (A) *Tbx1*, (B) *Flt1*, (C) *ApoE*, (D) *Btg1*, (E) *Tdo2*, (F) *Ecrg4*, and (G) *Rnf112* in stem/progenitor cells derived from the hippocampus of P0 C57BL/6J pups. Box-and-whisker plots (median + quartiles with maximum and minimum values). Genes that were altered were determined by Mann–Whitney non-parametric tests, corrected for multiple comparisons with an FDR of 10% (\**P* < 0.05; \*\**P* < 0.01). RQ, Relative quantification. *Tbx1* mRNA, Control siRNA, *N* = 25; *Tbx1* siRNA, *N* = 26; Flt1 mRNA, Control siRNA, *N* = 16; *Tbx1* siRNA, *N* = 17; ApoE mRNA, Control siRNA, *N* = 14; *Tbx1* siRNA, *N* = 15; *Btg1* mRNA, Control siRNA, *N* = 14; *Tbx1* siRNA, *N* = 15; *Tdo2* mRNA, Control siRNA, *N* = 14; *Tbx1* siRNA, *N* = 15; *Ecrg4* mRNA, Control siRNA, *N* = 9; *Tbx1* siRNA, *N* = 9; *Rnf112* mRNA, Control siRNA, *N* = 9; *Tbx1* siRNA, *N* = 9.

**Figure S16.**
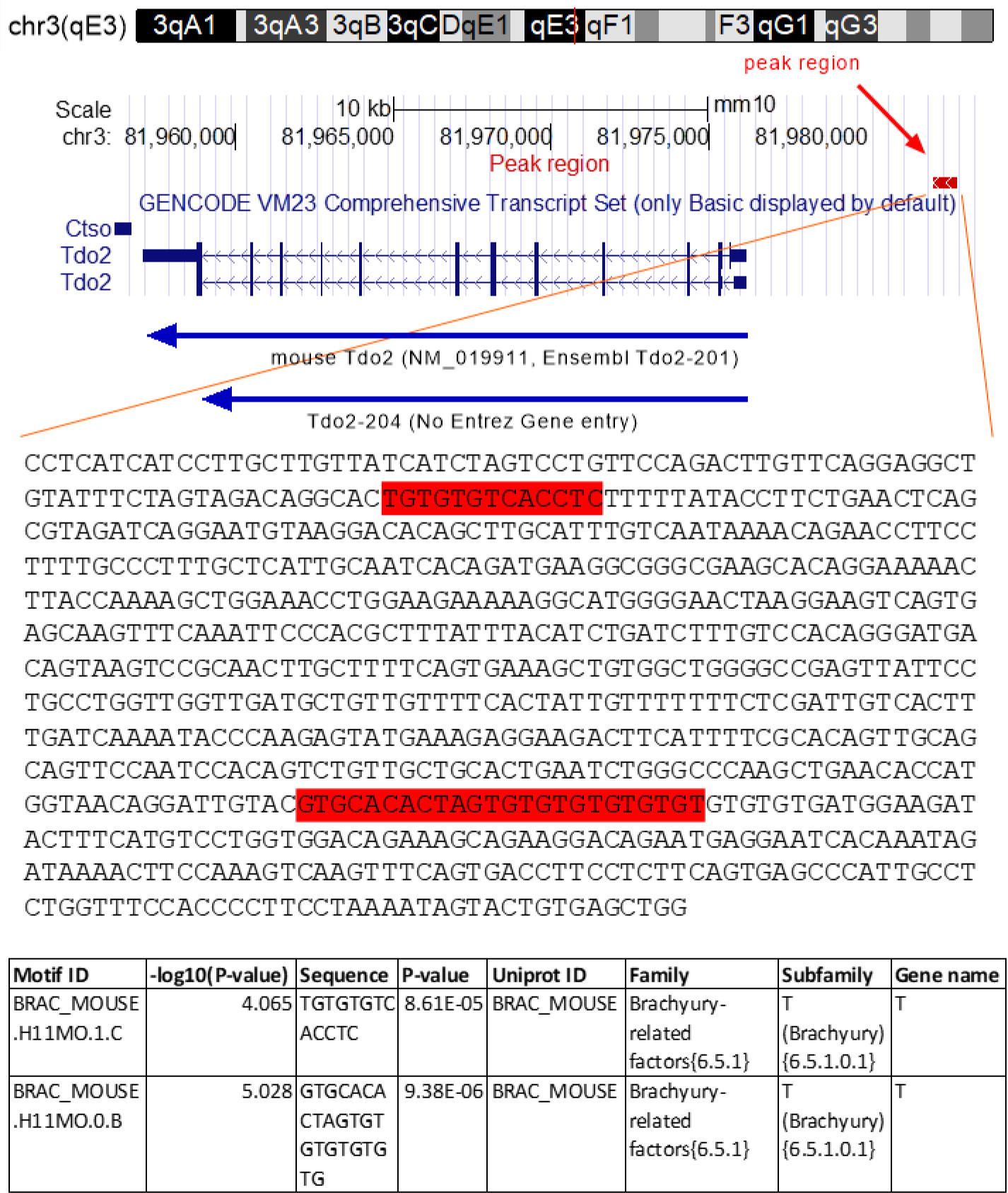
A peak representing the TBX1-binding site near *Tdo2* and its nucleotide sequences.

**Figure S17.**
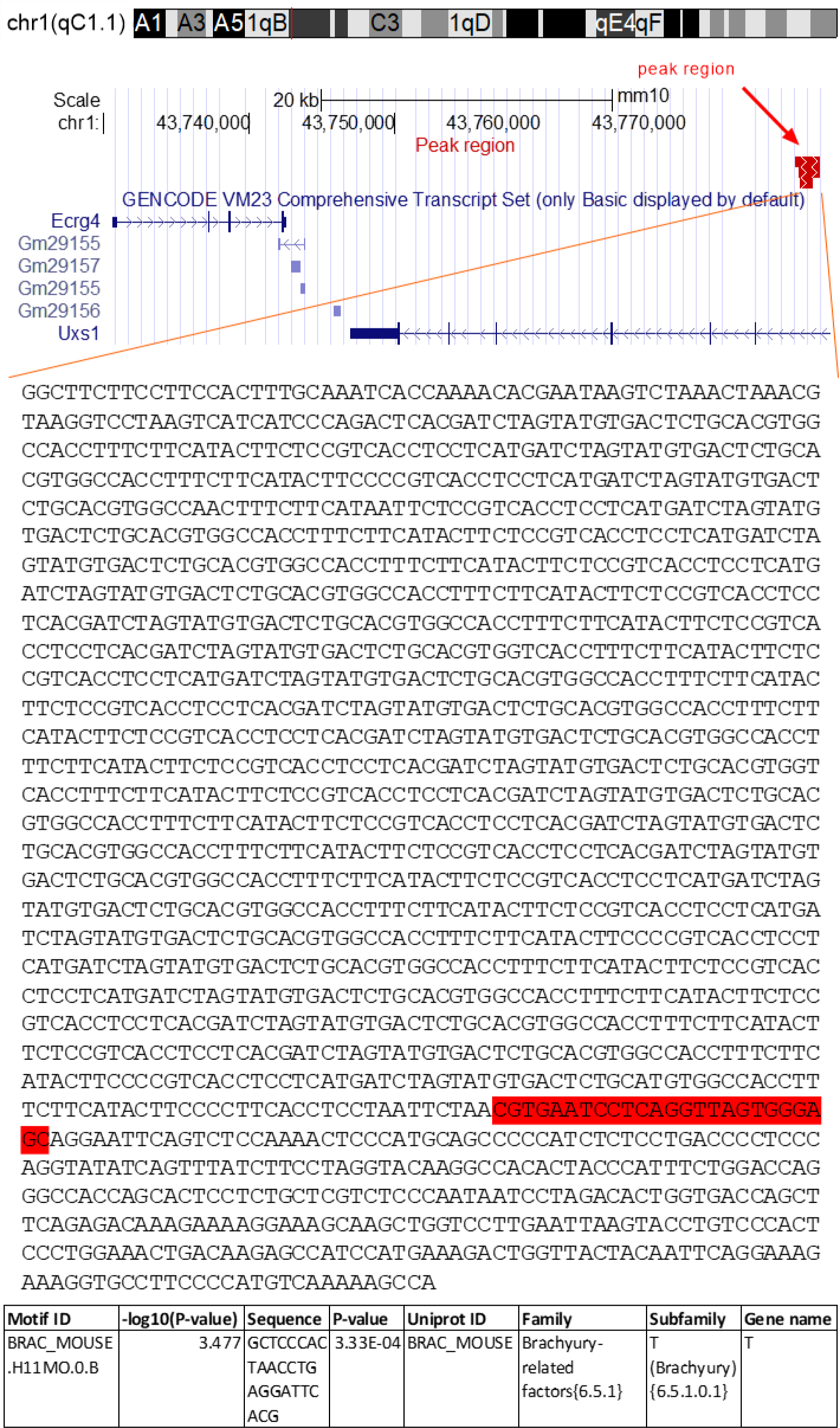
A peak representing the TBX1-binding site near *Ecrg4* and its nucleotide sequence.

**Figure S18.**
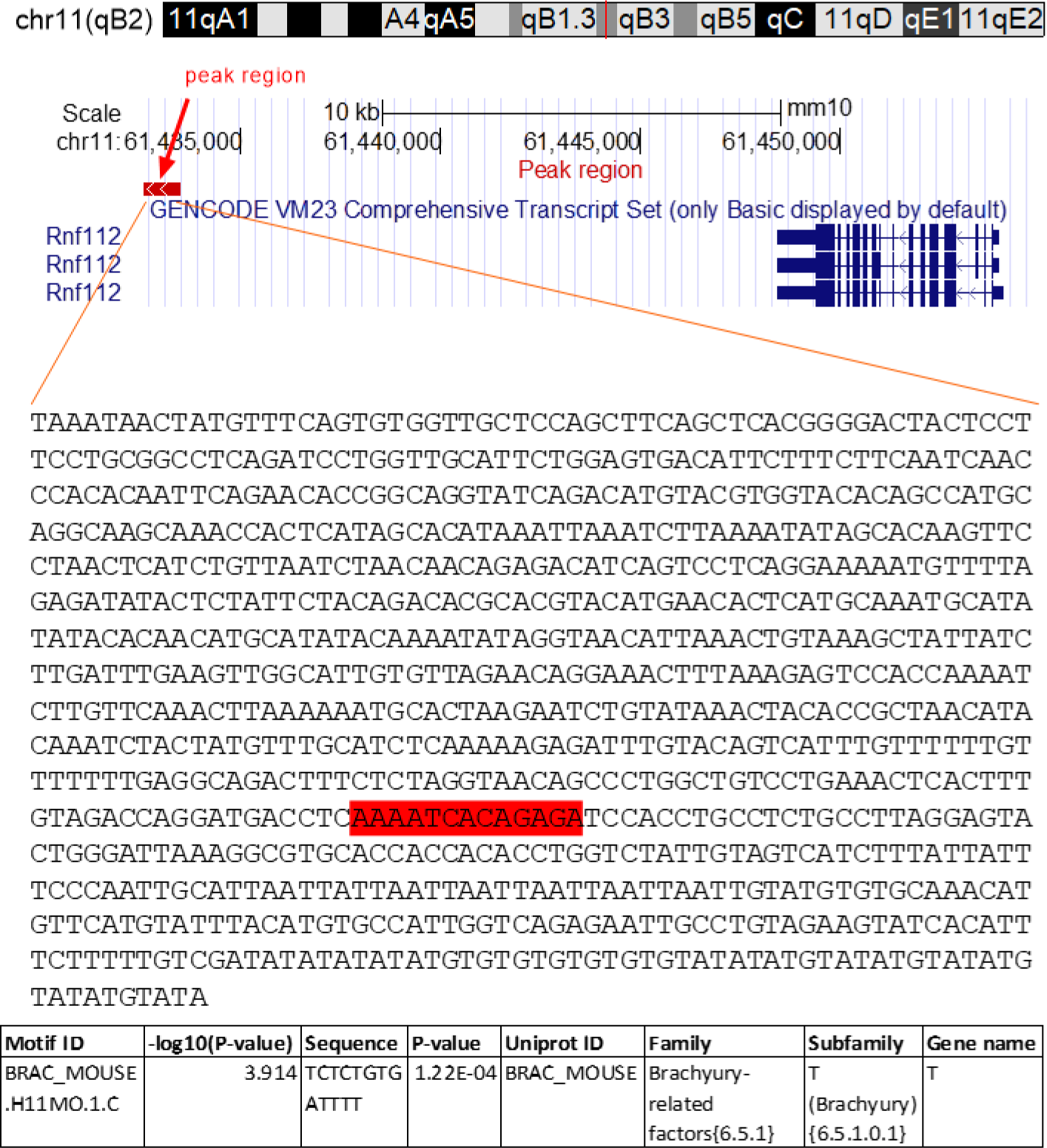
A peak representing the TBX1-binding site near *Rnf112* and its nucleotide sequence.

**Table S3.**
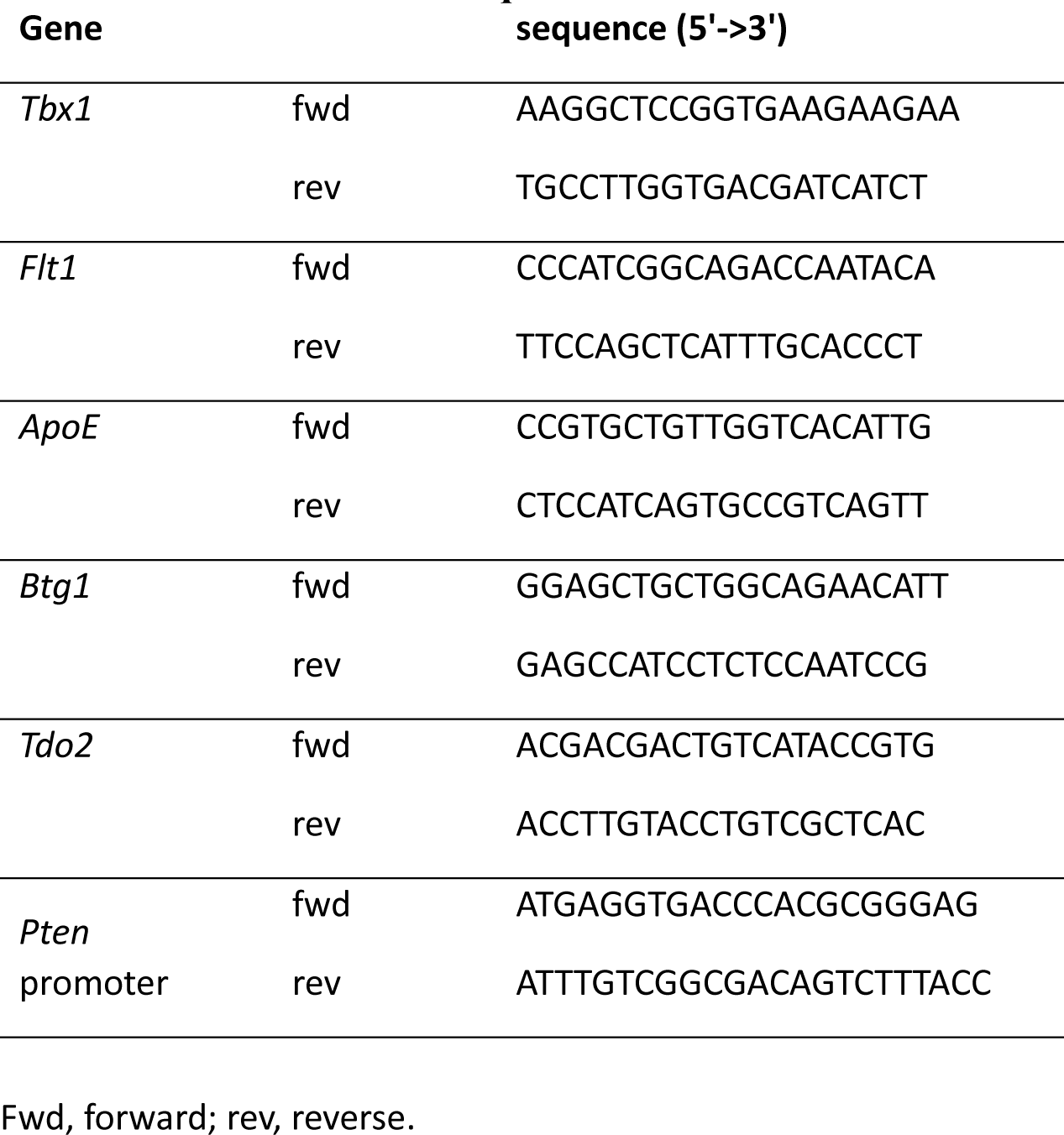
Primers used for qRT-PCR.

**Table S4.** List of antibodies used and their validation source.

| antibody | validated by | URL |
| --- | --- | --- |
| rat monoclonal BrdU antibody, BIO-RAD, Cat# OBT0030S, Hercules, CA) | vender | https://www.bio-rad-antibodies.com/static/datasheets/obt00/brdu-antibody-bu1-75-icr1-obt0030s.pdf |
| goat polyclonal DCX antibody (1:25, Santa Cruz, Cat# sc-8066, Dallas, TX) | other researchers | 225 citations at https://www.scbt.com/p/doublecortin-antibody-c-18?requestFrom=search |
| rabbit polyclonal GFP antibody (1:400, Thermo Fisher Scientific, Cat# A11122, Waltham, MA) | vender | verified at https://www.thermofisher.com/antibody/product/GFP-Antibody-Polyclonal/A-11122 |
| rabbit monoclonal TBX1 antibody (1:500, Abcam, Cat# ab109313, Cambridge, UK) | vender | https://www.abcam.com/tbx1-antibody-epr32892-ab109313.html |
| mouse monoclonal PTEN antibody (1:100, Santa Cruz, Cat# sc-7974, Dallas, TX) | vender | https://www.scbt.com/p/pten-antibody-a2b1?requestFrom=search |
| mouse monoclonal cyclin B1 antibody (1:100, Santa Cruz, Cat# sc-245, Dallas, TX) | vender | https://www.scbt.com/p/cyclin-b1-antibody-gns1?productCanUrl=cyclin-b1-antibody-gns1&_requestid=741230 |
| mouse monoclonal GAPDH antibody (1:500, Santa Cruz, Santa Cruz, Cat# sc-32233, Dallas, TX) | vender | https://www.scbt.com/p/gapdh-antibody-6c5?requestFrom=search |
| purified mouse monoclonal NeuN antibody, clone A60 (1:200, cat# MAB377, Millipore) | vender | https://www.emdmillipore.com/US/en/product/Anti-NeuN-Antibody-clone-A60,MM_NF-MAB377?ReferrerURL=https%3A%2F%2Fwww.google.com%2F&bd=1 |
| purified mouse Nestin antibody, Clone 25 (1:25, 1:50, ad 1:200, BD611658, BD Transduction Laboratories) | vender | https://www.bdbiosciences.com/en-us/products/reagents/western-blotting-and-molecular-reagents/western-blot-reagents/purified-mouse-anti-nestin.611658 |
| Donkey anti Rabbit IgG<br>Alexa-488 (1:200)<br>A21206 | vender | https://www.thermofisher.com/antibody/product/Donkey-anti-Rabbit-IgG-H-L-Highly-Cross-Adsorbed-Secondary-Antibody-Polyclonal/A-21206 |
| Donkey anti-Goat IgG<br>Alexa-594 (1:200)<br>A11058 | vender | https://www.thermofisher.com/antibody/product/Donkey-anti-Goat-IgG-H-L-Cross-Adsorbed-Secondary-Antibody-Polyclonal/A-11058 |
| Donkey anti-Mouse IgG<br>Alexa-594 (1:200)<br>A21203 | vender | https://www.thermofisher.com/antibody/product/Donkey-anti-Mouse-IgG-H-L-Highly-Cross-Adsorbed-Secondary-Antibody-Polyclonal/A-21203 |

**Table S5.** Reagents used and their validation.

| Reagent | Purity | Reference |
| --- | --- | --- |
| Tamoxifen (T5648; Sigma, St. Louis, MO, USA) | ≥99% | https://www.sigmaaldrich.com/US/en/product/sigma/t5648 |
| epidermal growth factor (EGF) (#236-EG-200; R&D systems, Minneapolis, MN, USA) | >97%, by SDS-PAGE visualized with Silver Staining and quantitative densitometry by Coomassie® Blue Staining | https://www.rndsystems.com/products/recombinant-human-egf-protein-cf_236-eg |
| Fibronectin (#1030-FN-05M, R&D) | >90%, by SDS-PAGE under reducing conditions and visualized by silver stain | https://www.rndsystems.com/products/bovine-fibronectin-protein-cf_1030-fn |
| BrdU (10 µM, 1 h, B9285; Sigma) | ≥99% | https://www.sigmaaldrich.com/US/en/product/sigma/b9285 |
| DAPI (D3571; Invitrogen, Waltham, MA, USA) | HPLC ≥96% at 254 nm | https://www.thermofisher.com/order/catalog/product/D3571#/D3571 |
| Thymidine (T1895-5G; Sigma) | ≥99% | https://www.sigmaaldrich.com/US/en/product/sigma/t1895 |

## References and Notes

1. T. Singh, B.M. Neale, M.J Daly, SCHEMA, Exome sequencing identifies rare coding variants in 10 genes which confer substantial risk for schizophrenia. MedRxiv, (2020).

2. F. K. Satterstrom, J. A. Kosmicki, J. Wang, M. S. Breen, S. De Rubeis, J. Y. An, M. Peng, R. Collins, J. Grove, L. Klei, C. Stevens, J. Reichert, M. S. Mulhern, M. Artomov, S. Gerges, B. Sheppard, X. Xu, A. Bhaduri, U. Norman, H. Brand, G. Schwartz, R. Nguyen, E. E. Guerrero, C. Dias, Consortium Autism Sequencing, Psych-Broad Consortium i, C. Betancur, E. H. Cook, L. Gallagher, M. Gill, J. S. Sutcliffe, A. Thurm, M. E. Zwick, A. D. Borglum, M. W. State, A. E. Cicek, M. E. Talkowski, D. J. Cutler, B. Devlin, S. J. Sanders, K. Roeder, M. J. Daly, J. D. Buxbaum, Large-Scale Exome Sequencing Study Implicates Both Developmental and Functional Changes in the Neurobiology of Autism. Cell 180, 568–584 e523 (2020).

3. I. Mitra, B. Huang, N. Mousavi, N. Ma, M. Lamkin, R. Yanicky, S. Shleizer-Burko, K. E. Lohmueller, M. Gymrek, Patterns of de novo tandem repeat mutations and their role in autism. Nature 589, 246–250 (2021).

4. J. Zinkstok, E. Boot, A.S. Bassett, N. Hiroi, N.J. Butcher, C. Vingerhoets, J.A.S. Vorstman, T.A.M.J. van Amelsvoort, The 22q11.2 deletion syndrome from a neurobiological perspective. Lancet Psychiatry 6, 951–960. (2019).

5. T. Ogata, T. Niihori, N. Tanaka, M. Kawai, T. Nagashima, R. Funayama, K. Nakayama, S. Nakashima, F. Kato, M. Fukami, Y. Aoki, Y. Matsubara, TBX1 mutation identified by exome sequencing in a Japanese family with 22q11.2 deletion syndrome-like craniofacial features and hypocalcemia. PLoS. One 9, e91598 (2014).

6. R. Paylor, B. Glaser, A. Mupo, P. Ataliotis, C. Spencer, A. Sobotka, C. Sparks, C.H. Choi, J. Oghalai, S. Curran, K.C. Murphy, S. Monks, N. Williams, M.C. O’Donovan, M.J. Owen, P.J. Scambler, E. Lindsay, Tbx1 haploinsufficiency is linked to behavioral disorders in mice and humans: implications for 22q11 deletion syndrome. Proc. Natl. Acad. Sci. U. S. A 103, 7729–7734 (2006).

7. T. R. Insel, B. N. Cuthbert, Medicine. Brain disorders? Precisely. Science 348, 499–500 (2015).

8. R.E. Gur, J.J. Yi, D.M. Donald-McGinn, S.X. Tang, M.E. Calkins, D. Whinna, M.C. Souders, Savitt, E.H. Zackai, P.J. Moberg, B.S. Emanuel, R.C. Gur, Neurocognitive development in 22q11.2 deletion syndrome: comparison with youth having developmental delay and medical comorbidities. Mol. Psychiatry 19, 1205–1211 (2014).

9. H. Stefansson, A. Meyer-Lindenberg, S. Steinberg, B. Magnusdottir, K. Morgen, S. Arnarsdottir, G. Bjornsdottir, G. B. Walters, G. A. Jonsdottir, O. M. Doyle, H. Tost, O. Grimm, S. Kristjansdottir, H. Snorrason, S. R. Davidsdottir, L. J. Gudmundsson, G. F. Jonsson, B. Stefansdottir, I. Helgadottir, M. Haraldsson, B. Jonsdottir, J. H. Thygesen, A. J. Schwarz, M. Didriksen, T. B. Stensbol, M. Brammer, S. Kapur, J. G. Halldorsson, S. Hreidarsson, E. Saemundsen, E. Sigurdsson, K. Stefansson, CNVs conferring risk of autism or schizophrenia affect cognition in controls. Nature 505, 361–366 (2014).

10. N. Hiroi, T. Yamauchi, Modeling and Predicting Developmental Trajectories of Neuropsychiatric Dimensions Associated With Copy Number Variations. Int J Neuropsychopharmacol 22, 488–500 (2019).

11. T. Hiramoto, G. Kang, G. Suzuki, Y. Satoh, R. Kucherlapati, Y. Watanabe, N. Hiroi, Tbx1: identification of a 22q11.2 gene as a risk factor for autism spectrum disorder in a mouse model. Hum Mol Genet 20, 4775–4785 (2011).

12. R. Kato, A. Machida, K. Nomoto, G. Kang, T. Hiramoto, K. Tanigaki, K. Mogi, N. Hiroi, T. Kikusui, Maternal approach behaviors toward neonatal calls are impaired by mother’s experiences of raising pups with a risk gene variant for autism. Dev Psychobiol 63, 108–113 (2021).

13. T. Takahashi, S. Okabe, P. O. Broin, A. Nishi, K. Ye, M. V. Beckert, T. Izumi, A. Machida, G. Kang, S. Abe, J. L. Pena, A. Golden, T. Kikusui, N. Hiroi, Structure and function of neonatal social communication in a genetic mouse model of autism. Mol Psychiatry 21, 1208–1214 (2016).

14. M. J. Hawrylycz, E. S. Lein, A. L. Guillozet-Bongaarts, E. H. Shen, L. Ng, J. A. Miller, L. N. van de Lagemaat, K. A. Smith, A. Ebbert, Z. L. Riley, C. Abajian, C. F. Beckmann, A. Bernard, D. Bertagnolli, A. F. Boe, P. M. Cartagena, M. M. Chakravarty, M. Chapin, J. Chong, R. A. Dalley, B. David Daly, C. Dang, S. Datta, N. Dee, T. A. Dolbeare, V. Faber, D. Feng, D. R. Fowler, J. Goldy, B. W. Gregor, Z. Haradon, D. R. Haynor, J. G. Hohmann, S. Horvath, R. E. Howard, A. Jeromin, J. M. Jochim, M. Kinnunen, C. Lau, E. T. Lazarz, C. Lee, T. A. Lemon, L. Li, Y. Li, J. A. Morris, C. C. Overly, P. D. Parker, S. E. Parry, M. Reding, J. J. Royall, J. Schulkin, P. A. Sequeira, C. R. Slaughterbeck, S. C. Smith, A. J. Sodt, S. M. Sunkin, B. E. Swanson, M. P. Vawter, D. Williams, P. Wohnoutka, H. R. Zielke, D. H. Geschwind, P. R. Hof, S. M. Smith, C. Koch, S. G. N. Grant, A. R. Jones, An anatomically comprehensive atlas of the adult human brain transcriptome. Nature 489, 391–399 (2012).

15. D.W. Meechan, T.M. Maynard, Y. Wu, D. Gopalakrishna, J.A. Lieberman, A.S. LaMantia, Gene dosage in the developing and adult brain in a mouse model of 22q11 deletion syndrome. Mol. Cell Neurosci 33, 412–428 (2006).

16. G. Flore, S. Cioffi, M. Bilio, E. Illingworth, Cortical Development Requires Mesodermal Expression of Tbx1, a Gene Haploinsufficient in 22q11.2 Deletion Syndrome. Cereb Cortex 27, 2210–2225 (2017).

17. D. W. Meechan, E. S. Tucker, T. M. Maynard, A. S. LaMantia, Diminished dosage of 22q11 genes disrupts neurogenesis and cortical development in a mouse model of 22q11 deletion/DiGeorge syndrome. Proc Natl Acad Sci U S A 106, 16434–16445 (2009).

18. S.A. Bayer, Development of the hippocampal region in the rat. I. Neurogenesis examined with 3H-thymidine autoradiography. J. Comp Neurol 190, 87–114 (1980).

19. A.J. Eisch, C.D. Mandyam, Adult neurogenesis: can analysis of cell cycle proteins move us “Beyond BrdU”? Curr. Pharm. Biotechnol 8, 147–165 (2007).

20. P. Taupin, BrdU immunohistochemistry for studying adult neurogenesis: paradigms, pitfalls, limitations, and validation. Brain Res Rev 53, 198–214 (2007).

21. C. J. Bostock, D. M. Prescott, J. B. Kirkpatrick, An evaluation of the double thymidine block for synchronizing mammalian cells at the G1-S border. Exp Cell Res 68, 163–168 (1971).

22. C. A. Denny, N. S. Burghardt, D. M. Schachter, R. Hen, M. R. Drew, 4- to 6-week-old adult-born hippocampal neurons influence novelty-evoked exploration and contextual fear conditioning. Hippocampus 22, 1188–1201 (2012).

23. G. S. Kirshenbaum, S. R. Lieberman, T. J. Briner, E. D. Leonardo, A. Dranovsky, Adolescent but not adult-born neurons are critical for susceptibility to chronic social defeat. Front Behav Neurosci 8, 289 (2014).

24. L. Garrett, J. Zhang, A. Zimprich, K. M. Niedermeier, H. Fuchs, V. Gailus-Durner, M. Hrabe de Angelis, D. Vogt Weisenhorn, W. Wurst, S. M. Holter, Conditional Reduction of Adult Born Doublecortin-Positive Neurons Reversibly Impairs Selective Behaviors. Front Behav Neurosci 9, 302 (2015).

25. A. R. Pereira-Caixeta, L. O. Guarnieri, D. C. Medeiros, Emam Mendes, L. C. D. Ladeira, M. T. Pereira, M. F. D. Moraes, G. S. Pereira, Inhibiting constitutive neurogenesis compromises long-term social recognition memory. Neurobiol Learn Mem 155, 92–103 (2018).

26. A. C. von Mering, L. J. Jensen, B. Snel, S. D. Hooper, M. Krupp, M. Foglierini, N. Jouffre, M. Huynen, P. Bork, STRING: known and predicted protein-protein associations, integrated and transferred across organisms. Nucleic Acids Res 33, D433–437 (2005).

27. B. M. Gasperini, A. J. Hill, J. L. McFaline-Figueroa, B. Martin, S. Kim, M. D. Zhang, D. Jackson, Leith, J. Schreiber, W. S. Noble, C. Trapnell, N. Ahituv, J. Shendure, A Genome-wide Framework for Mapping Gene Regulation via Cellular Genetic Screens. Cell 176, 377–390 e319 (2019).

28. S. M. Lloyd, X. Bao, Pinpointing the Genomic Localizations of Chromatin-Associated Proteins: The Yesterday, Today, and Tomorrow of ChIP-seq. Curr Protoc Cell Biol 84, e89 (2019).

29. G. Broitman-Maduro, M. Owraghi, W. W. Hung, S. Kuntz, P. W. Sternberg, M. F. Maduro, The NK-2 class homeodomain factor CEH-51 and the T-box factor TBX-35 have overlapping function in C. elegans mesoderm development. Development 136, 2735–2746 (2009).

30. A. Amiri, W. Cho, J. Zhou, S.G. Birnbaum, C.M. Sinton, R.M. McKay, L.F. Parada, Pten deletion in adult hippocampal neural stem/progenitor cells causes cellular abnormalities and alters neurogenesis. J. Neurosci 32, 5880–5890 (2012).

31. R. Savino, M. Carotenuto, A. N. Polito, S. Di Noia, M. Albenzio, A. Scarinci, A. Ambrosi, F. Sessa, N. Tartaglia, G. Messina, Analyzing the Potential Biological Determinants of Autism Spectrum Disorder: From Neuroinflammation to the Kynurenine Pathway. Brain Sci 10, (2020).

32. E. S. Lein, M. J. Hawrylycz, N. Ao, M. Ayres, A. Bensinger, A. Bernard, A. F. Boe, M. S. Boguski, K. S. Brockway, E. J. Byrnes, L. Chen, L. Chen, T. M. Chen, M. C. Chin, J. Chong, B. E. Crook, A. Czaplinska, C. N. Dang, S. Datta, N. R. Dee, A. L. Desaki, T. Desta, E. Diep, T. A. Dolbeare, M. J. Donelan, H. W. Dong, J. G. Dougherty, B. J. Duncan, A. J. Ebbert, G. Eichele, L. K. Estin, C. Faber, B. A. Facer, R. Fields, S. R. Fischer, T. P. Fliss, C. Frensley, S. N. Gates, K. J. Glattfelder, K. R. Halverson, M. R. Hart, J. G. Hohmann, M. P. Howell, D. P. Jeung, R. A. Johnson, P. T. Karr, R. Kawal, J. M. Kidney, R. H. Knapik, C. L. Kuan, J. H. Lake, A. R. Laramee, K. D. Larsen, C. Lau, T. A. Lemon, A. J. Liang, Y. Liu, L. T. Luong, J. Michaels, J. J. Morgan, R. J. Morgan, M. T. Mortrud, N. F. Mosqueda, L. L. Ng, R. Ng, G. J. Orta, C. C. Overly, T. H. Pak, S. E. Parry, S. D. Pathak, O. C. Pearson, R. B. Puchalski, Z. L. Riley, H. R. Rockett, S. A. Rowland, J. J. Royall, M. J. Ruiz, N. R. Sarno, K. Schaffnit, N. V. Shapovalova, T. Sivisay, C. R. Slaughterbeck, S. C. Smith, K. A. Smith, B. I. Smith, A. J. Sodt, N. N. Stewart, K. R. Stumpf, S. M. Sunkin, M. Sutram, A. Tam, C. D. Teemer, C. Thaller, C. L. Thompson, L. R. Varnam, A. Visel, R. M. Whitlock, P. E. Wohnoutka, C. K. Wolkey, V. Y. Wong, M. Wood, M. B. Yaylaoglu, R. C. Young, B. L. Youngstrom, X. F. Yuan, B. Zhang, T. A. Zwingman, A. R. Jones, Genome-wide atlas of gene expression in the adult mouse brain. Nature 445, 168–176 (2007).

33. M. Kanai, H. Funakoshi, H. Takahashi, T. Hayakawa, S. Mizuno, K. Matsumoto, T. Nakamura, Tryptophan 2,3-dioxygenase is a key modulator of physiological neurogenesis and anxiety-related behavior in mice. Mol Brain 2, 8 (2009).

34. O. H. Yilmaz, R. Valdez, B. K. Theisen, W. Guo, D. O. Ferguson, H. Wu, S. J. Morrison, Pten dependence distinguishes haematopoietic stem cells from leukaemia-initiating cells. Nature 441, 475–482 (2006).

35. J. Zhang, J. C. Grindley, T. Yin, S. Jayasinghe, X. C. He, J. T. Ross, J. S. Haug, D. Rupp, K. S. Porter-Westpfahl, L. M. Wiedemann, H. Wu, L. Li, PTEN maintains haematopoietic stem cells and acts in lineage choice and leukaemia prevention. Nature 441, 518–522 (2006).

36. Y. Nakatani, H. Kiyonari, T. Kondo, Ecrg4 deficiency extends the replicative capacity of neural stem cells in a Foxg1-dependent manner. Development 146, (2019).

37. F. K. Satterstrom, R. K. Walters, T. Singh, E. M. Wigdor, F. Lescai, D. Demontis, J. A. Kosmicki, J. Grove, C. Stevens, J. Bybjerg-Grauholm, M. Baekvad-Hansen, D. S. Palmer, J. B. Maller, Psych-Broad Consortium i, M. Nordentoft, O. Mors, E. B. Robinson, D. M. Hougaard, T. M. Werge, P. Bo Mortensen, B. M. Neale, A. D. Borglum, M. J. Daly, Autism spectrum disorder and attention deficit hyperactivity disorder have a similar burden of rare protein-truncating variants. Nat Neurosci 22, 1961–1965 (2019).

38. P. F. Sullivan, D. H. Geschwind, Defining the Genetic, Genomic, Cellular, and Diagnostic Architectures of Psychiatric Disorders. Cell 177, 162–183 (2019).

39. W. Gong, S. Gottlieb, J. Collins, A. Blescia, H. Dietz, E. Goldmuntz, D.M. Donald-McGinn, E.H. Zackai, B.S. Emanuel, D.A. Driscoll, M.L. Budarf, Mutation analysis of TBX1 in non-deleted patients with features of DGS/VCFS or isolated cardiovascular defects. J. Med. Genet 38, E45 (2001).

40. K. Hasegawa, H. Tanaka, Y. Higuchi, Y. Hayashi, K. Kobayashi, H. Tsukahara, Novel heterozygous mutation in TBX1 in an infant with hypocalcemic seizures. Clin Pediatr Endocrinol 27, 159–164 (2018).

41. D. Szklarczyk, A. L. Gable, D. Lyon, A. Junge, S. Wyder, J. Huerta-Cepas, M. Simonovic, N. T. Doncheva, J. H. Morris, P. Bork, L. J. Jensen, C. V. Mering, STRING v11: protein-protein association networks with increased coverage, supporting functional discovery in genome-wide experimental datasets. Nucleic Acids Res 47, D607–D613 (2019).

42. A. Sebe-Pedros, A. Ariza-Cosano, M. T. Weirauch, S. Leininger, A. Yang, G. Torruella, M. Adamski, M. Adamska, T. R. Hughes, J. L. Gomez-Skarmeta, I. Ruiz-Trillo, Early evolution of the T-box transcription factor family. Proc Natl Acad Sci U S A 110, 16050–16055 (2013).

43. J.L. Mignone, V. Kukekov, A.S. Chiang, D. Steindler, G. Enikolopov, Neural stem and progenitor cells in nestin-GFP transgenic mice. J. Comp Neurol 469, 311–324 (2004).

44. D.C. Lagace, M.C. Whitman, M.A. Noonan, J.L. Ables, N.A. DeCarolis, A.A. Arguello, M.H. Donovan, S.J. Fischer, L.A. Farnbauch, R.D. Beech, R.J. DiLeone, C.A. Greer, C.D. Mandyam, A.J. Eisch, Dynamic contribution of nestin-expressing stem cells to adult neurogenesis. J. Neurosci 27, 12623–12629 (2007).

45. J.S. Arnold, E.M. Braunstein, T. Ohyama, A.K. Groves, J.C. Adams, M.C. Brown, B.E. Morrow, Tissue-specific roles of Tbx1 in the development of the outer, middle and inner ear, defective in 22q11DS patients. Hum. Mol. Genet 15, 1629–1639 (2006).

46. J. Battiste, A.W. Helms, E.J. Kim, T.K. Savage, D.C. Lagace, C.D. Mandyam, A.J. Eisch, G. Miyoshi, J.E. Johnson, Ascl1 defines sequentially generated lineage-restricted neuronal and oligodendrocyte precursor cells in the spinal cord. Development 134, 285–293 (2007).

47. D. Petrik, S. Yun, S. E. Latchney, S. Kamrudin, J. A. LeBlanc, J. A. Bibb, A. J. Eisch, Early postnatal in vivo gliogenesis from nestin-lineage progenitors requires cdk5. PLoS One 8, e72819 (2013).

48. A. M. Ceasrine, N. Ruiz-Otero, E. E. Lin, D. N. Lumelsky, E. D. Boehm, R. Kuruvilla, Tamoxifen Improves Glucose Tolerance in a Delivery-, Sex-, and Strain-Dependent Manner in Mice. Endocrinology 160, 782–790 (2019).

49. M.Y. Sun, M.J. Yetman, T.C. Lee, Y. Chen, J.L. Jankowsky, Specificity and efficiency of reporter expression in adult neural progenitors vary substantially among nestin-CreER(T2) lines. J. Comp Neurol 522, 1191–1208 (2014).

50. S. Boku, T. Izumi, S. Abe, T. Takahashi, A. Nishi, H. Nomaru, Y. Naka, G. Kang, M. Nagashima, A. Hishimoto, S. Enomoto, G. Duran-Torres, K. Tanigaki, J. Zhang, K. Ye, S. Kato, P.T. Mannisto, K. Kobayashi, N. Hiroi, Copy number elevation of 22q11.2 genes arrests the developmental maturation of working memory capacity and adult neurogenesis. Molecular Psychiatry 23, 985–992 (2018).

51. M. Toritsuka, S. Kimoto, K. Muraki, M.A. Landek-Salgado, A. Yoshida, N. Yamamoto, Y. Horiuchi, H. Hiyama, K. Tajinda, N. Keni, E. Illingworth, T. Iwamoto, T. Kishimoto, A. Sawa, K. Tanigaki, Deficits in microRNA-mediated Cxcr4/Cxcl12 signaling in neurodevelopmental deficits in a 22q11 deletion syndrome mouse model. Proc. Natl. Acad. Sci. U. S. A 110, 17552–17557 (2013).

52. H. A. Cameron, R. D. McKay, Adult neurogenesis produces a large pool of new granule cells in the dentate gyrus. J Comp Neurol 435, 406–417 (2001).

53. M. Nakamura, K. Ye, M. Barbachan E Silva, T. Yamauchi, D. Hoeppner, A. Fayyazuddin, G. Kang, E. Yuda, M. Nagashima, S. Enomoto, T. Hiramoto, R. Sharp, I. Kaneko, K. Tajinda, M. Adachi, T. Mihara, S. Tokuno, M. Geyer, P. O’Broin, M. Matsumoto, N. Hiroi, Computational identification of variables in neonatal vocalizations predictive for post-pubertal social behaviors in a mouse model of 16p11.2 deletion. Molecular Psychiatry Online Published on April 15, 2021, (2021).

54. C. Y. McLean, D. Bristor, M. Hiller, S. L. Clarke, B. T. Schaar, C. B. Lowe, A. M. Wenger, G. Bejerano, GREAT improves functional interpretation of cis-regulatory regions. Nat Biotechnol 28, 495–501 (2010).

55. W. J. Kent, C. W. Sugnet, T. S. Furey, K. M. Roskin, T. H. Pringle, A. M. Zahler, D. Haussler, The human genome browser at UCSC. Genome Res 12, 996–1006 (2002).

56. I. V. Kulakovskiy, I. E. Vorontsov, I. S. Yevshin, R. N. Sharipov, A. D. Fedorova, E. I. Rumynskiy, Y. A. Medvedeva, A. Magana-Mora, V. B. Bajic, D. A. Papatsenko, F. A. Kolpakov, V. J. Makeev, HOCOMOCO: towards a complete collection of transcription factor binding models for human and mouse via large-scale ChIP-Seq analysis. Nucleic Acids Res 46, D252–D259 (2018).

57. B. Rhead, D. Karolchik, R. M. Kuhn, A. S. Hinrichs, A. S. Zweig, P. A. Fujita, M. Diekhans, K. E. Smith, K. R. Rosenbloom, B. J. Raney, A. Pohl, M. Pheasant, L. R. Meyer, K. Learned, F. Hsu, J. Hillman-Jackson, R. A. Harte, B. Giardine, T. R. Dreszer, H. Clawson, G. P. Barber, D. Haussler, W. J. Kent, The UCSC Genome Browser database: update 2010. Nucleic Acids Res 38, D613–619 (2010).

58. G. Yu, L. G. Wang, Q. Y. He, ChIPseeker: an R/Bioconductor package for ChIP peak annotation, comparison and visualization. Bioinformatics 31, 2382–2383 (2015).

59. S. Durinck, Y. Moreau, A. Kasprzyk, S. Davis, B. De Moor, A. Brazma, W. Huber, BioMart and Bioconductor: a powerful link between biological databases and microarray data analysis. Bioinformatics 21, 3439–3440 (2005).

60. Richard Iannone. (2020), pp. https://rich-iannone.github.io/DiagrammeR/.

